## Supplementary Information for "Demultiplexing and barcode-specific adaptive sampling for nanopore direct RNA sequencing"

### Supplementary Tables

| Data set | Sequencing run | RNA template | Replicate | RTA barcode | RNA barcode |
|---|---|---|---|---|---|
| 1 | 1 | *V. chol.* glycine riboswitch | 1 | DPC_1 | bc13 |
| | | | | DPC_2 | bc14 |
| | | | | DPC_3 | bc15 |
| | | | | DPC_4 | bc16 |
| 2 | 2 | Tetrahymena ribozyme | 2 | DPC_1 | bc04 |
| | | | | DPC_2 | bc05 |
| | | | | DPC_3 | bc17 |
| | | | | DPC_4 | bc18 |
| 3 | 3 | *V. chol.* glycine riboswitch | 1 | WDX_bc01 | bc12 |
| | | | | WDX_bc02 | bc04 |
| | | | | WDX_bc03 | bc05 |
| | | | | WDX_bc04 | bc03 |
| | | | | WDX_bc05 | bc06 |
| | | | | WDX_bc06 | bc07 |
| | | | | WDX_bc07 | bc01 |
| | | | | WDX_bc08 | bc09 |
| | | | | WDX_bc09 | bc02 |
| | | | | WDX_bc10 | bc10 |
| | | | | WDX_bc11 | bc11 |
| | | | | WDX_bc12 | bc08 |
| 4 | 3 | hc16 | 2 | WDX_bc01 | bc10 |
| | | | | WDX_bc02 | bc09 |
| | | | | WDX_bc03 | bc01 |
| | | | | WDX_bc04 | bc08 |
| | | | | WDX_bc05 | bc05 |
| | | | | WDX_bc06 | bc07 |
| | | | | WDX_bc07 | bc11 |
| | | | | WDX_bc08 | bc03 |
| | | | | WDX_bc09 | bc06 |
| | | | | WDX_bc10 | bc04 |
| | | | | WDX_bc11 | bc02 |
| | | | | WDX_bc12 | bc12 |
| 5 | 4 | HCV IRES | 3 | WDX_bc01 | bc05 |
| | | | | WDX_bc02 | bc06 |
| | | | | WDX_bc03 | bc07 |
| | | | | WDX_bc04 | bc01 |
| | | | | WDX_bc05 | bc11 |
| | | | | WDX_bc06 | bc03 |
| | | | | WDX_bc07 | bc04 |
| | | | | WDX_bc08 | bc10 |
| | | | | WDX_bc09 | bc02 |
| | | | | WDX_bc10 | bc09 |
| | | | | WDX_bc11 | bc08 |
| | | | | WDX_bc12 | bc12 |
| 6 | 4 | bact. RNase P (not poly-adenylated) | 4 | WDX_bc01 | bc09 |
| | | | | WDX_bc02 | bc11 |
| | | | | WDX_bc03 | bc10 |
| | | | | WDX_bc04 | bc08 |
| | | | | WDX_bc05 | bc12 |
| | | | | WDX_bc06 | bc04 |
| | | | | WDX_bc07 | bc01 |
| | | | | WDX_bc08 | bc07 |
| | | | | WDX_bc09 | bc02 |
| | | | | WDX_bc10 | bc05 |
| | | | | WDX_bc11 | bc03 |
| | | | | WDX_bc12 | bc06 |

**Table S1:** Run-RNA-barcode to RTA assignments for RTA training/evaluation data

| DPC-RTA | **Replicate1** *V. chol* gly Polyadenylated | **Replicate2** tetrahymena Polyadenylated | Total |
|---|---|---|---|
| Barcode 1 | 601 | 1856 | 2457 |
| Barcode 2 | 600 | 2482 | 3082 |
| Barcode 3 | 191 | 3141 | 3332 |
| Barcode 4 | 558 | 975 | 1533 |
| Total | 1950 | 8454 | 10404 |

**Table S2:** Generated experimental DPC-RTA data, coverage per barcode and replicate after mapping and adapter detection.

| WDX-RTA | **Replicate1** *V. chol* gly Polyadenylated | **Replicate2** hc16 Polyadenylated | **Replicate3** HCV IRES Polyadenylated | **Replicate4** bact RNase P Not polyadenylated | Total |
|---|---|---|---|---|---|
| Barcode 01 | 548 | 2604 | 411 | 872 | 4435 |
| Barcode 02 | 633 | 3493 | 712 | 1476 | 6314 |
| Barcode 03 | 433 | 3125 | 868 | 894 | 5320 |
| Barcode 04 | 652 | 3078 | 188 | 1182 | 5100 |
| Barcode 05 | 111 | 2132 | 237 | 285 | 2765 |
| Barcode 06 | 162 | 963 | 626 | 1480 | 3231 |
| Barcode 07 | 888 | 4806 | 701 | 1207 | 7602 |
| Barcode 08 | 3120 | 2962 | 703 | 1894 | 8679 |
| Barcode 09 | 324 | 5839 | 427 | 1305 | 7895 |
| Barcode 10 | 1006 | 2802 | 857 | 2073 | 6738 |
| Barcode 11 | 588 | 2459 | 888 | 1649 | 5584 |
| Barcode 12 | 570 | 5575 | 360 | 749 | 7254 |
| Total | 9035 | 39838 | 6978 | 15066 | 70917 |

**Table S3:** Generated experimental WDX-RTA data, coverarge per barcode and replicate after mapping and adapter detection.

| Barcode ID | Barcode sequence |
|---|---|
| Barcode 1 | GGCTTCTTCTTGCTCTTAGG |
| Barcode 2 | GTGATTCTCGTCTTTCTGCG |
| Barcode 3 | GTACTTTTCTCTTTGCGCGG |
| Barcode 4 | GGTCTTCGCTCGGTCTTATT |

**Table S4:** DeePlexiCon RTA barcodes, sequences in 5’ to 3’ direction [7].

| Barcode ID | Barcode sequence |
|---|---|
| Barcode 1 | TTTTTACTGCCAGTGACT |
| Barcode 2 | AGGGGAGAGAGCCCCCCC |
| Barcode 3 | CACGTCATTTTCCACGTC |
| Barcode 4 | GGAGGCCAGGCGGACCGA |
| Barcode 5 | ACGGACCTTTTGACTTAA |
| Barcode 6 | TATTGCATACTGCGCCGC |
| Barcode 7 | CCACGGAGGGAGGATTGG |
| Barcode 8 | TTACCGGCAGTGACGGAC |
| Barcode 9 | CGAGATTGCATCCCCCCC |
| Barcode 10 | TACCACCTGCCGGCGGCC |
| Barcode 11 | GCCCGCCGGGGGAGAAGC |
| Barcode 12 | TTTTTTTTACCGGCAGTT |

**Table S5:** Optimized WarpDemuX barcodes, sequences in 5‘ to 3‘ direction.

### Supplementary Methods

#### SM.1 Designing WarpDemuX Barcodes

##### Experimental design

The RTA adapter consists of two partially complementary oligos. To ensure compatibility with the sequencing chemistry we designed the adapter oligos similarly to the tested design described in Smith et al. [7], with the exception of allowing variation of GC content. In detail, the two last nucleotides of the 5' end of the forward RTA adapter (GG) were kept constant to reduce potential ligation biases. In addition, all adapters retained the 3' adapter sequence that is required for the RMX motor protein to ligate onto. The reverse adapter was designed as reverse complement of the forward adapter, with the exception of a 10 nucleotide dT 3' overhang to hybridize to the RNA poly(A) tail, as well as a Y anchor sequence 5' of the reverse complement RTA barcode. All oligos were ordered from IDT with standard desalting purification, and with forward adapters containing a 5' phosphorylation. The following scheme exemplifies the design:

| | |
|---|---|
| forward adapter | /5Phos/GG–[*WDX barcode*]–GGTAGTAGGTTC |
| reverse adapter | GAGGCGAGCGGTCAATTTT–[*revcomp–WDX barcode*]–CCTTTTTTTTTT |

**Table SM.1.1:** Custom RTA design scheme (5' to 3' orientation for both oligos)

##### *In silico* design

Barcode optimization addresses the need for a set of distinguishable signals that can be easily detected and separated. For barcodes of length $L$, an exhaustive search across all possible sets of $k$ barcodes has complexity $\binom{4^L}{k}$, making such a search computationally infeasible: $\binom{6.9\times10^{10}}{12} \approx 2.3 \times 10^{121}$ for $L = 18$, $k = 12$ .

Instead, we constructed a set of wave-like target signal patterns since we expect such patterns to improve segmentation, as well as distinction between barcodes and thus make for good barcodes in the context of our classification method.

###### Target Patterns Derivation

Across target patterns, we combined wave-like elements with diverse frequencies and amplitudes. Additionally, we maintained consistent minimal and maximal values across all target patterns to mitigate potential normalization biases during signal processing.

The customizable section of the RTA that serves as barcode spans 18 nucleotides. Given the sequencing k-mer size of 6, however, the preceding five constant nucleotides and the following constant two, as well as the first bases of the subsequent polyA tail also influence its signal. As a result, we expect 20 DNA and 3 DNA-polyA signal events. As there is no ONT k-mer model for mixed DNA-polyA k-mers, we assumed the signal of these k-mers to be an interpolation between the value of the last DNA k-mer and the expected signal level of a pure polA segment ($\sim$ 110 pA). When constructing the target patterns, we substituted the interpolation k-mers with a singular polyA segment observation, thus yielding patterns that span 21 observations.

###### Candidate Set Construction

We searched for barcode sequences whose putative signal, constructed using the ONT k-mer models (`github.com/nanoporetech/kmer_models`), would closely match these target patterns. These models are designed for DNA sequenced in the 5' to 3' direction. In dRNA-seq, however, the RTA is sequenced in 3' to 5' direction. To increase the likelihood that our modeled barcode signals reflect the sequencing data, we limited our barcode design to the subset of k-mers for which the 5'-3' to 3'-5' directional reversal likely has little effect. Specifically, we selected k-mers with minimal variation (less than 10 pA) in expected signal mean when compared to their mirrored counterparts. For example, if the expected signal level difference between the k-mer AAACCC and the k-mer CCCAAA is $<$ 10 pA, then the k-mer AAACCC is selected.

Subsequently, we discretized the expected signal values for the selected k-mers (1524 in total) and their mirrored counterparts respectively into five equally-sized, rank-based buckets. Only those k-mers whose original and mirrored versions both fell within the lowest, middle, or highest rank buckets were further considered, leaving us with 531 k-mers. This discretization and selection strategy enhances the probability of identifying wave-like patterns–marked by alternating low and high values–when assembling the k-mers into barcodes. Additionally, this approach strengthens the stringency of our heuristic for ensuring robustness against directional reversal.

From the filtered set of k-mers, we generated all conceivable combinations of five k-mers, adhering to a stride of three. This means that the final three bases of one k-mer overlap with the initial three bases of the following k-mer. This approach yielded roughly $3.4 \times 10^6$ candidate barcode sequences, each 18 bases long.

For each, we constructed the full adapter sequence by pre- and appending the flanking regions of adapter. Using the ONT k-mer models we constructed the putative adapter signal and appended it with a singular observation of the expected polyA tail signal. The full adapter signal was normalized to zero mean and unit variance. We then selected the last 21 observations of the idealized adapter signal, corresponding to the variable barcode part and calculated the distance to each target pattern.

#### Candidate Selection

To select the set of $k = 12$ barcodes we selected the top 50 best-scoring (lowest distance) candidate barcodes per target pattern. For the $k = 12$ target patterns highlighted in Fig. 2A, this yielded a total of 600 unique barcodes, for which we computed all $600^2$ pairwise DTWDs.

Lastly, within the matrix of $600^2$ pairwise DTWDs we greedily approximate the Maximal Diversity Problem (MDP) to propose 12 candidate barcodes that we expect to be accurately identifiable and distinguishable. The greedy heuristic used to approximate the MDP was implemented as in Algorithm SM.1.1.

**Algorithm SM.1.1:** Greedy heuristic for solving the Maximum Diversity Problem.

```
input  : Distance matrix D ∈ ℝ^{n×n}_{≥0}, Number of elements to select m
output: Indices of the selected elements
n ← number of rows in D;
first ← max_i(∑_j D_ij); // Select the element with the maximum sum of distances
selected ← [first];
while length of selected < m do
    maxMinDist ← −1;
    nextIndex ← −1;
    for i ← 0 to n − 1 do
        if i ∉ selected then
            minDist2Selected ← min{D_ij : j ∈ selected};
            if minDist2Selected > maxMinDist then
                maxMinDist ← minDist2Selected;
                nextIndex ← i;
            end
        end
    end
    Append nextIndex to selected;
end
return selected;
```

#### SM.2 WarpDemuX Signal Processing

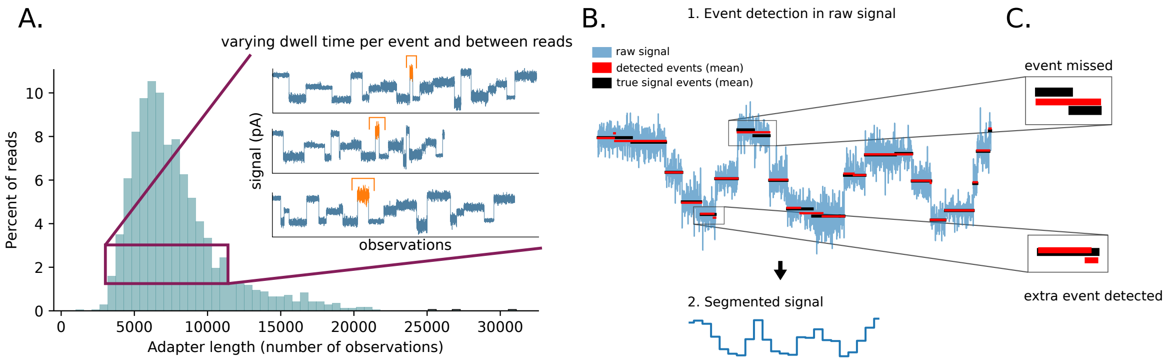


**Figure SM.2.1:** Intrinsic variation in dwell time poses a source of noise. (A) Varying dwell time per event and between reads results in varying length adapter signals. (B) Segmentation of the raw signal into events, with each segment represented by its average signal, reduces signal noise caused by differences in dwell time and within-event measurement noise (C) Signal noise causes imperfect segmentation in which some events are missed while other spurious events are detected.

To account for the inherent variation in dwell time (Fig. SM.2.1A), the raw signal is segmented (Fig. SM.2.1B) by identifying the most significant shifts in current level based on the running difference between neighboring windows of raw signal. The pseudo code is described in SM.2.1. Its parameters were tuned to reflect the translocation speed of the RTA, with a minimal segment length of 15 observations a sliding window width of 30 observations. Considering the known length of the adapter sequence, the signal is expected to contain 98 events. However, because segmentation is likely to be imperfect (Fig. SM.2.1C), with missed events posing a greater challenge than spurious events, we intentionally oversegment the signal by approximately 10% to detect 110 events. Each segmented event is then represented by its average signal. Post-segmentation, segment means are normalized to zero mean and unit variance. Only the variable barcode component of the segmented signal, expected to be represented within the last 25 segments, based on the known barcode sequence length and location in the adapter, is used for classification.

**Algorithm SM.2.1:** Greedy algorithm for raw signal segmentation based on the most significant shifts in signal intensity.

```
input : Raw signal X of length N
l ← 6; // Minimal segment length
k ← 12; // Width of sliding window
n ← 110; // Number of segmentation points to identify
W ← ∅; // Empty list
T ← ∅;
C ← [0, N];
S ← ∅;
for i ← 1 to N − k do
    w_i ← X[i : i + k];
    Append w_i to W;
end
for i ← 1 to N − 2k do
    w̄_i ← mean of w_i;
    σ_i^2 ← variance of w_i;
    w̄_{i+1} ← mean of w_{i+1};
    σ_{i+1}^2 ← variance of w_{i+1};
    t_{i,i+1} ← |w̄_{i+1} − w̄_i| / sqrt(σ_i^2 + σ_{i+1}^2);
    Append t_{i,i+1} to T;
end
while size of C < n do
    Find index i of max absolute value in T;
    Append i + k to C;
    Exclude values within range l of i from T;
end
Sort values in C;
for each consecutive pair (c_1, c_2) in C do
    s ← mean of X[c_1 : c_2];
    Append s to S;
end
return S;
```

Segmentation ensures that all inputs to the classification procedure are of uniform length. Due to the signal normalization, pairwise DTWDs are of similar magnitude across barcodes, negating the need for further scaling or normalization of the pairwise distance matrix.

### SM.3 WarpDemuX Parameterization

#### SVM

We use an C-Support Vector Classification SVM implementation based on libsvm [34], with a one-vs-one scheme for multi-class support, as implemented in Scikit-Learn [35] (scikit-learn 1.3.1).

With `C`, the regularization parameter, set to 10.0, `kernel = 'precomputed'` and default settings otherwise.

#### DTWD

To compute the Dynamic Time Warping Distance (DTWD) between two barcode fingerprints, we use the Python Package `dtaidistance 2.3.10` [32], with a window size of 15 (upper bound to allowed shift from the diagonal) and a additive warping penalty of 0.1 per compression or expansion.

### SM.4 Generating in vitro transcribed barcoded RNA

#### RNA barcoding strategy

In order to train and benchmark RTA-based barcoding strategies we generated in vitro transcribed RNAs of different templates containing varying RNA-encoded barcodes. This allows us to generate high accuracy ground truth labels for different RTA signals. To ensure that no signal spillover from RNA barcode into RTA signal was occuring we decided to enzymatically poly(A) tail these RNAs post in vitro transcription (although they already contain a 10 nt poly(A) tail that can be used for direct ligation of RTA adapters). Fig. SM.4.1 provides a visual summary of the protocol. Further, we also decided to randomize RNA-barcode to RTA assignments to eliminate the possibility of inadvertently training on RNA signal in the rare possibility of RTA adapter signal failure S1 .

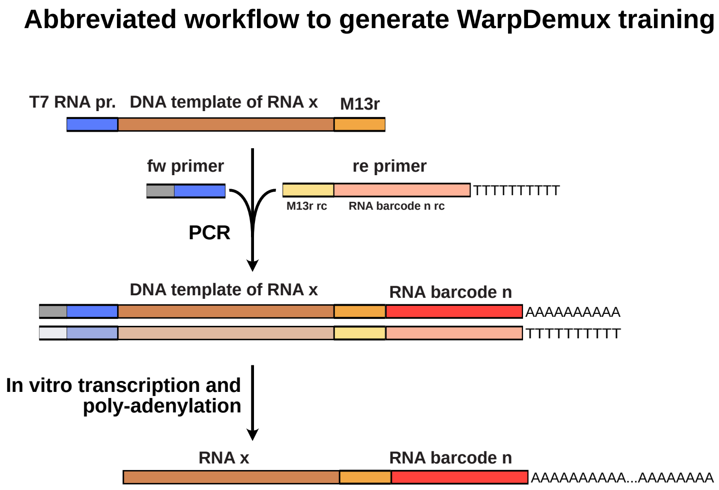


**Figure SM.4.1:** Abbreviated workflow of experimental protocol to generate WarpDemuX training data.

### RNA barcode design

The RNA barcode sequences were designed with high RNA sequencing error rates in mind. Specifically, a large pool of 8 nt long barcode candidates was generated by random sampling, with some additional candidates supplemented by BARCOSEL [39], and a set of 18 barcodes that maximized inter-barcode Levenshtein edit distances was chosen. To balance purine and pyrimidine content in each barcode and double the number of variable nucleotides in the final barcodes a self-complementary stem-loop (i.e. [barcode]-GAGUA-[revcomp_barcode]) was generated for each barcode. To introduce these RNA barcodes into DNA templates reverse primers containing a 10 nt long T stretch at their 5' end, followed by the reverse complement of the barcode stem-loop, and finally the reverse complement of the M13-reverse sequence were ordered with standard desalting purity from IDT (Table SM.4.1).

| RNA barcode ID | Barcode unique sequence | Reverse primer for PCR |
|---|---|---|
| RNA_bc01 | GATAGTGCgagtaGCACTATC | TTTTTTTTTTGATAGTGCTACTCGCACTATCcaggaaacagctatgaccatg |
| RNA_bc02 | GCAATACTgagtaAGTATTGC | TTTTTTTTTTGCAATACTTACTCAGTATTGCcaggaaacagctatgaccatg |
| RNA_bc03 | GATTATCAgagtaTGATAATC | TTTTTTTTTTGATTATCATACTCTGATAATCcaggaaacagctatgaccatg |
| RNA_bc04 | AGGTCAGGgagtaCCTGACCT | TTTTTTTTTTAGGTCAGGTACTCCCTGACCTcaggaaacagctatgaccatg |
| RNA_bc05 | GAGGCTCCgagtaGGAGCCTC | TTTTTTTTTTGAGGCTCCTACTCGGAGCCTCcaggaaacagctatgaccatg |
| RNA_bc06 | GCAACCAGgagtaCTGGTTGC | TTTTTTTTTTGCAACCAGTACTCCTGGTTGCcaggaaacagctatgaccatg |
| RNA_bc07 | GACTCCATgagtaATGGAGTC | TTTTTTTTTTGACTCCATTACTCATGGAGTCcaggaaacagctatgaccatg |
| RNA_bc08 | GTGTAATCgagtaGATTACAC | TTTTTTTTTTGTGTAATCTACTCGATTACACcaggaaacagctatgaccatg |
| RNA_bc09 | ATGACGAAgagtaTTCGTCAT | TTTTTTTTTTATGACGAATACTCTTCGTCATcaggaaacagctatgaccatg |
| RNA_bc10 | CATGTTGAgagtaTCAACATG | TTTTTTTTTTCATGTTGATACTCTCAACATGcaggaaacagctatgaccatg |
| RNA_bc11 | CGAACTTCgagtaGAAGTTCG | TTTTTTTTTTCGAACTTCTACTCGAAGTTCGcaggaaacagctatgaccatg |
| RNA_bc12 | TATCTGTGgagtaCACAGATA | TTTTTTTTTTTTGAGTACTACTCGTACTCAAcaggaaacagctatgaccatg |
| RNA_bc13 | GACGGACCgagtaGGTCCGTC | TTTTTTTTTTGACGGACCTACTCGGTCCGTCcaggaaacagctatgaccatg |
| RNA_bc14 | GAGGTCTGgagtaCAGACCTC | TTTTTTTTTTGAGGTCTGTACTCCAGACCTCcaggaaacagctatgaccatg |
| RNA_bc15 | GCTCAATCgagtaGATTGAGC | TTTTTTTTTTGCTCAATCTACTCGATTGAGCcaggaaacagctatgaccatg |
| RNA_bc16 | GACGATACgagtaGTATCGTC | TTTTTTTTTTGACGATACTACTCGTATCGTCcaggaaacagctatgaccatg |
| RNA_bc17 | GACCGCCTgagtaAGGCGGTC | TTTTTTTTTTGACCGCCTTACTCAGGCGGTCcaggaaacagctatgaccatg |
| RNA_bc18 | GAGTCCTTgagtaAAGGACTC | TTTTTTTTTTGAGTCCTTTACTCAAGGACTCcaggaaacagctatgaccatg |

**Table SM.4.1:** RNA barcode design and reverse PCR primers to generate in vitro transcription templates. The constant M13r sequence is in lowercase.

### PCR to generate barcoded DNA templates

PCR was performed on templates of the *Vibrio cholerae* glycine riboswitch, hc16 ligase, bacterial RNase P type A, tetrahymena riboswitch and Hepatitis C Virus IRES cloned into pJet (Thermo Fisher) flanked by a T7 RNA promoter sequence and an M13r sequence at 5' and 3' ends respectively as described in Bohn et al. [2] using Primestar GXL (Takara) according to the manufacturer's instructions. Reaction volume was 20 ul, annealing temp. 50 °C, extension time 30 seconds at 35 cycles with forward primers described in Table SM.4.2 and reverse primers as described in Table SM.4.1.

| Template | FW Primer Sequence |
|---|---|
| HCV IRES | AAAGAAGACTTGGGGTAATACGACTCACTATAGGCCAGCCCCCGATTG |
| *V. chol.* Glycine riboswitch | TTCTAATACGACTCACTATAGGTCCGTTGAAGACTGCAGGAGAGTGG |
| bact. Rnase P | TTCTAATACGACTCACTATAGGAGAGGAGCAGGC |
| hc16 | TTCTAATACGACTCACTATAGGTAGACTCGCAGGAAGTCTACCGAGTAAGAGAAAGAGGA |
| Tetrahymena ribozyme | TTCTAATACGACTCACTATAGGAGGGAAAAGTTATCAGGCATGCACCTGGT |

**Table SM.4.2:** Forward PCR primers to generate in vitro transcription templates.

### In vitro transcription

PCR products were quality controlled via agarose gel and purified by addition of 18 ul H2O and 38 ul SPRI beads (Omega Biotek NGS). After hybridization for 5 min under light agitation (300 rpm on hula mixer) beads were pelleted using a DynaMag-96 Side Magnet (Invitrogen). Supernatant was removed, beads were washed twice with 100 ul 80% EtOH and eluted in 20 ul H2O. DNA concentration was measured with Nanodrop and 250 fmol of DNA was then used in a 10 ul in vitro transcription reaction: 5 mM NTPs, 5 U/ul T7 RNA polymerase (homemade), 1 ul YIPP (NEB), 0.1 ul RNasin (Molox) in 40 mM Tris pH 7.5, 18 mM $MgCl_2$, 10 mM DTT, 1 mM spermidine, 5 mM NTPs, 40 U RNasin (Molox) and homemade T7 RNA polymerase. After in vitro transcription for 2 h at 37 °C the DNA template was digested by addition of 10 ul of DNase mix containing 1 ul 10x DNase I buffer and 0.2 ul of DNase I (NEB) followed by a 15 min incubation at 37 °C. The RNA was then purified by addition of 1.4 volumes of SPRI beads and two washes with 80% EtOH. Purified RNA was then quality controlled with a 1.5% agarose gel in 1x TAE and concentration was quantified via Nanodrop.

### Poly(A) tailing

For poly(A) tailing 1 ug of IVT RNA was incubated with 0.5 ul E. coli poly(A) Polymerase (NEB), 1 ul 10 mM ATP in 1x poly(A) polymerase reaction buffer in a 10 ul reaction for 30 min at 37 °C, followed by purification with 1.4 volumes of SPRI beads and two washes in 190 ul 80% EtOH before elution in 20 ul RNase-free H2O.

### SM.5 Direct RNA sequencing library preparation

#### Annealing of RTA adapters

The forward and reverse strands of the RTA oligos were annealed following a procedure similar to that described in Smith et al. [7]. Specifically, oligos were diluted to 10 uM each in 30 mM HEPES-KOH (pH 7.5), 100 mM K-Acetate, heated to 95 °C for 1 min and cooled to 25 °C at a rate of 0.5 °C/min. The annealed RTA adapters were then diluted to 1.4 uM with RNase-free H2O and stored at -20 °C until use.

#### RTA adapter ligation

Volumes were scaled for direct RNA sequencing on a Flongle flow cell according to the protocols.io protocol of Krause [40] 2020. In detail, for each sample, 100 ng of poly-adenylated in vitro transcribed RNA in 4.4 ul, was transferred into a PCR tube, followed by addition of 1 ul previously annealed custom RTA adapter, 1.6 ul 5x NEBNext Quick Ligation Buffer (NEB) and 1 ul T4 DNA Ligase high concentration (NEB). The reaction was mixed well by pipetting and incubated at room temperature for 20 min. To stop the ligation 2 ul of 0.5 M EDTA was added to each reaction, samples were pooled, purified with 0.8 volumes of SPRI beads washed twice with 80 % EtOH, and eluted in 20 ul RNase-free H2O.

#### Reverse Transcription

To perform the reverse transcription a master mix containing 1.9 ul H2O, 15.7 ul 3x MarathonRT buffer (150 mM Tris-HCl pH 8.3, 600 mM KCl, 60 % (v/v) glycerol), 2.35 ul 100 mM DTT, 0.2 ul RNasin (Molox), 1.9 ul 50 mM MgCl2, 3 ul of 10 mM dNTPs and 2 ul MarathonRT (produced in house as described in [2]) was prepared and added to the 20 ul RNA-RTA ligation. After incubation at 42 °C for 60 min the reaction was stopped by addition of 2 ul Proteinase K (NEB), followed by another incubation at 42 °C for 15 min. The reaction was purified with 1 volume SPRI beads, washed thrice with 190 ul 80 % EtOH, and eluted in 30 ul RNase-free water (volume was increased to adjust for higher total input amount due to multiplexing).

#### Motor protein ligation

To proceed with Nanopore sequencing we next ligated on the motor protein. For this we transferred 10 ul of purified reverse-transcribed RNA-RTA library into a 1.5 ml DNA LoBind tube (Eppendorf), followed by addition of 2.5 ul H2O, 4 ul 5x NEBNext Quick Ligation Buffer (NEB), 2 ul RMX motor protein adapter (ONT SQK-RNA002), and 1.5 ul T4 DNA Ligase high conc. (NEB). The reaction was mixed by pipetting and incubated for 20 min at RT, followed by purification with 20 ul SPRI beads, washed twice with RNA wash buffer (WSB, ONT SQK-RNA002). The library was then eluted in 9 ul elution buffer (EB, ONT SQK-RNA002) and 8 ul was transferred into a new 1.5 ml DNA LoBind tube. The library was then loaded onto a Flongle FLO-FLG001 flow cell for sequencing.

### SM.6 Data Acquisition and Basecalling software configuration

- MinKNOW `22.12.7`
- dorado `0.3.4`
- protocol `MIN106_RNA:FLO-FLG001:SQK-RNA002`
- dorado model: `rna002_70bps_hac@v3`

### Supplementary Notes

#### SN.1 Training set selection, mislabeled instances and Noise Class

We calculated the pairwise Dynamic Time Warping Distances (DTWDs) for all barcode fingerprint instances outside of validation or test sets. First, we identified potentially mislabeled instances based on the robust Z-score of their median within-class distance. Reads with an absolute value of at least the 98th percentile were excluded from the data set.

Upon inspection of the excluded data, we noticed that a substantial part of these reads were the result of a sequencing artifact known as a missed read split. The two common types were: (1) a concatenation of two complete reads, each consisting of an adapter, polyA and RNA signal (Fig. SN.1.1a), and (2) a concatenation of an 'adapter-only' read and a complete read, wherein the first consists of an adapter and subsequent polyA tail signal, but misses the RNA signal (Fig. SN.1.1b).

In both cases, the adapter signal of the first read in the read pair is detected, processed and classified. However, the ground truth label for this read, inferred based on a combination of the basecalled RNA transcript and RNA barcode, may belong to the second read of the concatenated pair, rather than the one processed.

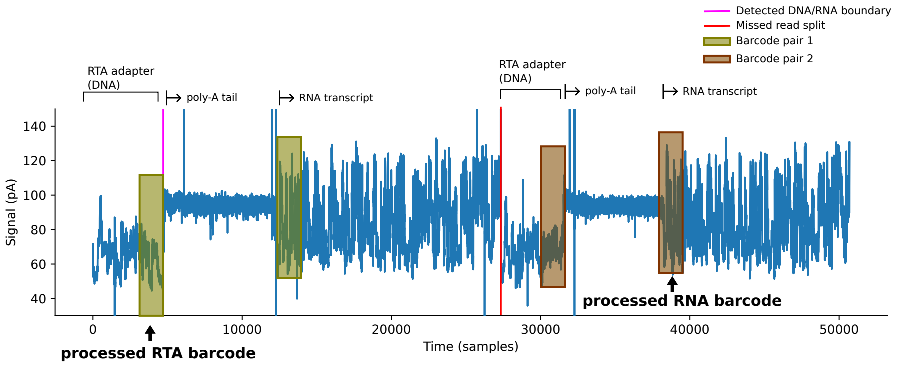


**(a)** Case 1

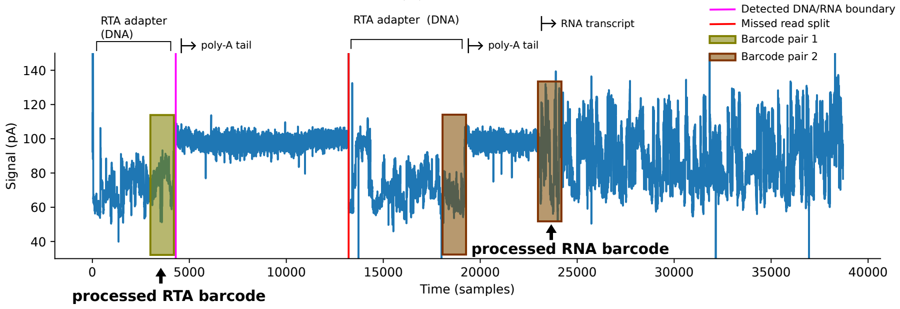


**(b)** Case 2

**Figure SN.1.1:** Examples of mislabeled signal instances due to missed read splits. (a) Concatenation of two complete reads, each consisting of an adapter, polyA and RNA signal. (b) Concatenation of an 'adapter-only' read and a complete read, wherein the first consists of an adapter and subsequent polyA tail signal, but misses the RNA signal.

After filtering out mislabeled instances, we calculated the robust Z-scores of the median inter-class distances to identify noisy signals. For this, we used the minimal absolute value score across classes per instance. Reads with a minimal absolute value score above the 99th percentile value were labeled as noise. From these, we randomly sampled 400 instances as training data for the noise class.

Upon inspection of the noise class data, we observed stalls to be the main type of signal irregularity, potentially leading to mis-segmentation of the adapter itself (Fig. SN.1.2a). In addition, we observed stalls in the beginning of the adapter to cause mis-identification of the DNA/RNA boundary (Fig. SN.1.2B).

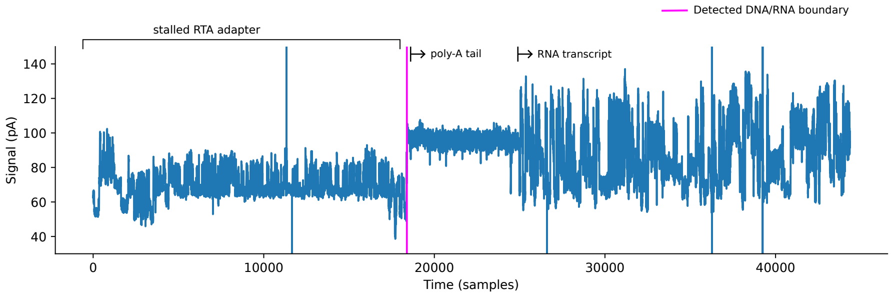


**(a)** Case 1

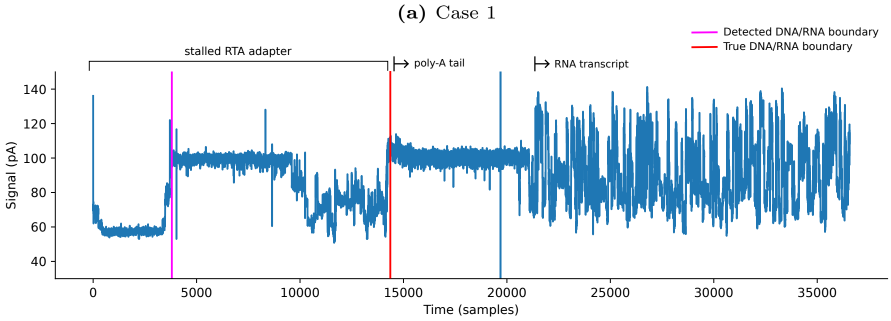


**(b)** Case 2

**Figure SN.1.2:** Examples of noise class signal instances. (a) Stall in adapter signal leading to potential mis-segmentation. (b) Stall in the beginning of the adapter leading to mis-identification of DNA/RNA boundary.

For the remaining reads, we approximate the solution of the Maximal- and Minimal Diversity Problem, stratified across barcode classes (Algorithm SN.1.1 and SN.1.2), to select 2x80 instances per barcode that respectively maximized and minimized their mutual distance. The training set was completed with 240 randomly sampled instances per barcode.

**Algorithm SN.1.1:** Greedy heuristic for solving the Maximum Diversity Problem, stratified over classes.

```
input  : Distance matrix D ∈ R_{≥0}^{n×n}, Number of elements to select m
output: Indices of the selected elements
n ← number of rows in D;
C ← Class labels of elements in D (rows);
selected ← []; // Empty list
for c ∈ unique(C) do
    first ← max_i(∑_j D_ij) : j ∈ selected AND i ∈ {C_i == c}; // Select the element with the maximum sum
        of distances per class
    Append first to selected;
end
while length of selected < m do
    for c ∈ unique(C) do
        maxMinDist ← −1;
        nextIndex ← −1;
        for i ← 0 to n − 1 do
            if i ∉ selected AND C_i == c then
                minDist2Selected ← min{D_ij : j ∈ selected};
                if minDist2Selected > maxMinDist then
                    maxMinDist ← minDist2Selected;
                    nextIndex ← i;
                end
            end
        end
        Append nextIndex to selected;
    end
end
return selected;
```

**Algorithm SN.1.2:** Greedy heuristic for solving the Minimum Diversity Problem, stratified over classes.

```
input  : Distance matrix D ∈ R_{≥0}^{n×n}, Number of elements to select m
output: Indices of the selected elements
n ← number of rows in D;
C ← Class labels of elements in D (rows);
selected ← []; // Empty list
for c ∈ unique(C) do
    first ← min_i(∑_j D_ij) : j ∈ selected AND i ∈ {C_i == c}; // Select the element with the minimum sum
        of distances per class
    Append first to selected;
end
while length of selected < m do
    for c ∈ unique(C) do
        minMaxDist ← 1000;
        nextIndex ← −1;
        for i ← 0 to n − 1 do
            if i ∉ selected AND C_i == c then
                maxDist2Selected ← max{D_ij : j ∈ selected};
                if maxDist2Selected < minMaxDist then
                    minMaxDist ← maxDist2Selected;
                    nextIndex ← i;
                end
            end
        end
        Append nextIndex to selected;
    end
end
return selected;
```

#### Number of training instances per barcode

During WarpDemuX training, we selected the number of training instances per barcode based on the average 10-fold cross-validated model performance on the validation set. The model reached optimal performance at a small number of training instances per barcode (Fig. SN.1.3a), while run time kept increasing with the number of training instances as expected (Fig. SN.1.3b). The model's performance was found to be robust with as few as 400 training instances per barcode.

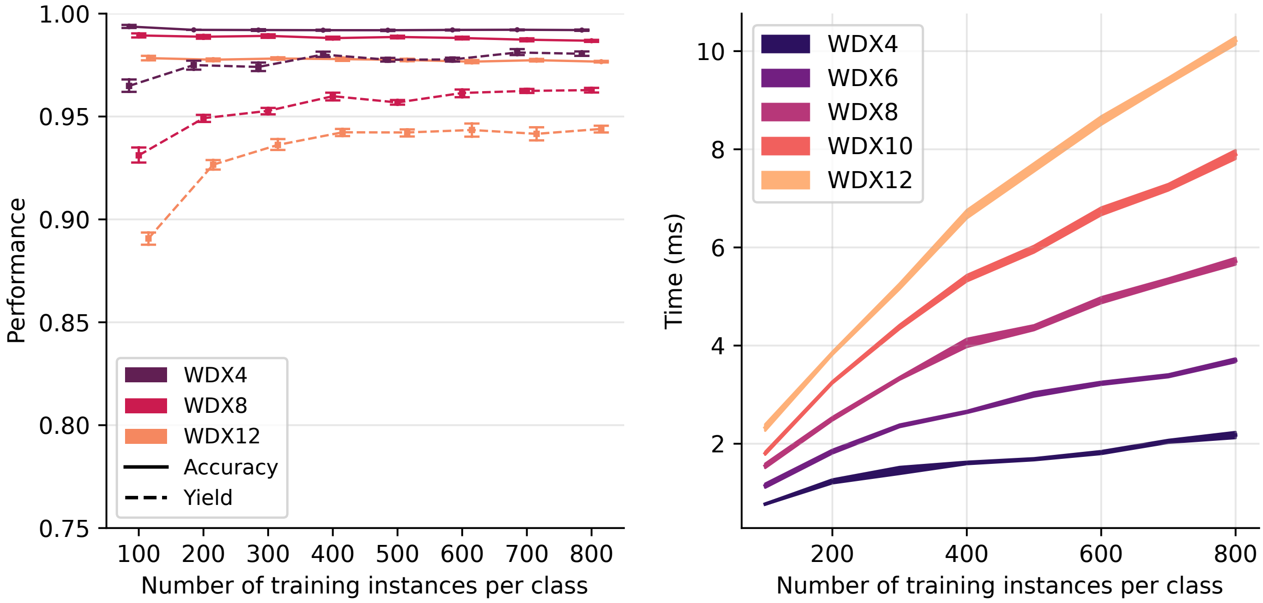


**Figure SN.1.3:** Performance on validation set during model selection. (a) Performance and yield for increasing number of training instances per barcode, evaluated at confidence threshold of 0.5, shows that the gains of increasing number of training instances per barcode on the model's quality already dimish at small training set sizes. Values represent mean over 10-fold cross-validation with errorbars indicating 95% CI. (b) Classification time per read for increasing number of training instances per barcode. Shaded area reflects the average ±2std classification time per read, computed from 10 runs of 4000 reads on 8 CPU cores (11th Gen Intel(R) Core(TM) i7-1165G7 (2.80GHz) with 32GB memory).

### SN.2 WarpDemuX performance

#### Best performing barcode sets

Based on a 10-fold cross-validated benchmark, we selected the best performing subsets of barcodes for experimental settings requiring less than 12 barcodes (Suppl. Table SN.2.1). The trained WarpDemuX models for these barcode subsets are available in the code repository.

| Model | Number of samples | WarpDemuX barcodes |
|---|---|---|
| WDX4 | 4 | 4 5 6 8 |
| WDX6 | 6 | 1 2 5 6 7 11 |
| WDX8 | 8 | 1 3 5 6 7 9 11 12 |
| WDX10 | 10 | 1 2 3 5 6 7 9 10 11 12 |
| WDX12 | 12 | 1 2 3 4 5 6 7 8 9 10 11 12 |

**Table SN.2.1:** Best performing barcodes sets for various numbers of barcodes, based on model selection benchmark.

#### Runtime

| Model | Time [ms] |
|---|---|
| WDX4 | 1.61 |
| WDX6 | 2.65 |
| WDX8 | 4.05 |
| WDX10 | 5.37 |
| WDX12 | 6.68 |

**Table SN.2.2:** Classification time per read for WarpDemuX models. Reported times reflect average time per read, computed from 10 runs of 4000 reads on 8 CPU cores (11th Gen Intel(R) Core(TM) i7-1165G7 (2.80GHz) with 32GB memory).

| Model | Time per read [ms] | |
|---|---|---|
| | mean | std |
| WDX4 | 14.37 | 0.07 |
| WDX6 | 14.62 | 0.12 |
| WDX8 | 14.93 | 0.07 |
| WDX10 | 15.26 | 0.09 |
| WDX12 | 15.56 | 0.14 |

**Table SN.2.3:** Run time per read for WarpDemuX models. Reported times reflect average time per read, computed from 10 runs of 4000 reads on 8 CPU cores (11th Gen Intel(R) Core(TM) i7-1165G7 (2.80GHz) with 32GB memory).

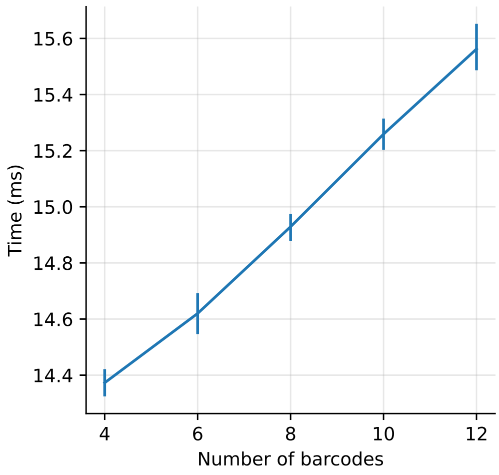


**Figure SN.2.1:** Run time per read for WarpDemuX models increases linearly with the number of barcodes. Reported times reflect average time per read, computed from 10 runs of 4000 reads on 8 CPU cores (11th Gen Intel(R) Core(TM) i7-1165G7 (2.80GHz) with 32GB memory).

#### Test set performances at different confidence thresholds

| Model | Dataset | Reads classified as noise [%] | Confidence cutoff | Unclassfied reads [%] | Accuracy | Precision | Recall |
|---|---|---|---|---|---|---|---|
| DPC | 1 | 0.000 | 0.000 | 0.000 | 0.873 | 0.860 | 0.853 |
| DPC | 1 | 0.000 | 0.300 | 6.634 | 0.905 | 0.896 | 0.886 |
| DPC | 1 | 0.000 | 0.500 | 11.020 | 0.925 | 0.919 | 0.907 |
| DPC | 1 | 0.000 | 0.800 | 19.751 | 0.956 | 0.952 | 0.944 |
| DPC | 2 | 0.000 | 0.000 | 0.000 | 0.833 | 0.785 | 0.846 |
| DPC | 2 | 0.000 | 0.300 | 9.164 | 0.878 | 0.831 | 0.891 |
| DPC | 2 | 0.000 | 0.500 | 15.916 | 0.906 | 0.859 | 0.918 |
| DPC | 2 | 0.000 | 0.800 | 26.206 | 0.954 | 0.916 | 0.963 |
| WDX-DPC | 1 | 0.000 | 0.000 | 0.000 | 0.932 | 0.915 | 0.938 |
| WDX-DPC | 1 | 0.000 | 0.300 | 5.179 | 0.951 | 0.939 | 0.954 |
| WDX-DPC | 1 | 0.000 | 0.500 | 9.487 | 0.965 | 0.957 | 0.967 |
| WDX-DPC | 1 | 0.000 | 0.800 | 18.462 | 0.982 | 0.977 | 0.983 |
| WDX-DPC | 2 | 0.000 | 0.000 | 0.000 | 0.937 | 0.929 | 0.919 |
| WDX-DPC | 2 | 0.000 | 0.300 | 4.093 | 0.957 | 0.950 | 0.944 |
| WDX-DPC | 2 | 0.000 | 0.500 | 7.275 | 0.969 | 0.964 | 0.958 |
| WDX-DPC | 2 | 0.000 | 0.800 | 14.443 | 0.985 | 0.980 | 0.979 |
| WDX4 | 3-5 | 1.750 | 0.000 | 0.000 | 0.984 | 0.984 | 0.984 |
| WDX4 | 3-5 | 1.750 | 0.300 | 1.375 | 0.987 | 0.987 | 0.987 |
| WDX4 | 3-5 | 1.750 | 0.500 | 2.708 | 0.989 | 0.989 | 0.989 |
| WDX4 | 3-5 | 1.750 | 0.800 | 6.083 | 0.992 | 0.992 | 0.992 |
| WDX4 | 6 | 4.668 | 0.000 | 0.000 | 0.978 | 0.964 | 0.974 |
| WDX4 | 6 | 4.668 | 0.300 | 2.541 | 0.984 | 0.978 | 0.980 |
| WDX4 | 6 | 4.668 | 0.500 | 4.648 | 0.987 | 0.983 | 0.984 |
| WDX4 | 6 | 4.668 | 0.800 | 9.977 | 0.990 | 0.990 | 0.988 |
| WDX12 | 3-5 | 2.167 | 0.000 | 0.000 | 0.960 | 0.961 | 0.960 |
| WDX12 | 3-5 | 2.167 | 0.300 | 3.750 | 0.976 | 0.976 | 0.976 |
| WDX12 | 3-5 | 2.167 | 0.500 | 6.181 | 0.980 | 0.980 | 0.980 |
| WDX12 | 3-5 | 2.167 | 0.800 | 13.444 | 0.986 | 0.986 | 0.986 |
| WDX12 | 6 | 4.752 | 0.000 | 0.000 | 0.941 | 0.937 | 0.943 |
| WDX12 | 6 | 4.752 | 0.300 | 6.000 | 0.965 | 0.963 | 0.965 |
| WDX12 | 6 | 4.752 | 0.500 | 10.056 | 0.975 | 0.973 | 0.975 |
| WDX12 | 6 | 4.752 | 0.800 | 19.010 | 0.984 | 0.983 | 0.983 |

**Table SN.2.4:** Comparative performance of model and barcode types at various confidence thresholds. Precision and recall as macro average across classes.

Stringent confidence cutoffs can be beneficial, for example in highly unbalanced samples in which the underrepresented sample is of interest.

| Model | Dataset | Reads classified as noise [%] | Confidence cutoff | Unclassfied reads [%] | Accuracy | Precision | Recall |
|---|---|---|---|---|---|---|---|
| DPC | 1 | 0.000 | 0.950 | 30.606 | 0.978 | 0.977 | 0.972 |
| DPC | 2 | 0.000 | 0.950 | 36.013 | 0.967 | 0.937 | 0.974 |
| WDX-DPC | 1 | 0.000 | 0.950 | 33.744 | 0.993 | 0.992 | 0.993 |
| WDX-DPC | 2 | 0.000 | 0.950 | 25.124 | 0.994 | 0.992 | 0.991 |
| WDX4 | 3-5 (test set) | 1.750 | 0.950 | 13.333 | 0.995 | 0.995 | 0.995 |
| WDX4 | 6 | 4.668 | 0.950 | 19.335 | 0.993 | 0.993 | 0.991 |
| WDX12 | 3-5 (test set) | 2.167 | 0.950 | 31.194 | 0.991 | 0.990 | 0.991 |
| WDX12 | 6 | 4.752 | 0.950 | 37.694 | 0.989 | 0.988 | 0.988 |

**Table SN.2.5:** Comparative performance of model and barcode types at stringent confidence thresholds 0.95. Precision and recall as macro average across classes.

| Model | Dataset | Reads classified as noise [%] | Confidence cutoff | Unclassfied reads [%] | Accuracy | Precision | Recall |
|---|---|---|---|---|---|---|---|
| DPC | 1 | 0.000 | 0.990 | 43.270 | 0.989 | 0.987 | 0.986 |
| DPC | 2 | 0.000 | 0.990 | 48.392 | 0.981 | 0.952 | 0.985 |
| WDX-DPC | 1 | 0.000 | 0.990 | 50.564 | 0.994 | 0.994 | 0.991 |
| WDX-DPC | 2 | 0.000 | 0.990 | 40.040 | 0.997 | 0.998 | 0.995 |
| WDX4 | 3-5 (test set) | 1.750 | 0.990 | 21.792 | 0.995 | 0.994 | 0.995 |
| WDX4 | 6 | 4.668 | 0.990 | 27.825 | 0.994 | 0.994 | 0.995 |
| WDX12 | 3-5 (test set) | 2.167 | 0.990 | 50.056 | 0.995 | 0.993 | 0.995 |
| WDX12 | 6 | 4.752 | 0.990 | 54.500 | 0.989 | 0.986 | 0.988 |

**Table SN.2.6:** Comparative performance of model and barcode types at stringent confidence thresholds 0.99. Precision and recall as macro average across classes.

#### Recommended confidence cutoffs

To facilitate a fair comparison between models with a different number of barcodes, as well as to improve the ease-of-use for the user, we introduce two modes of operation per model: a High Accuracy mode (HA) and a High Recovery mode (HR), with respectively tuned confidence cutoffs based on the performance on the validation set.

For the mode HA, the confidence cutoff is set to the value for a 90% yield on the validation set. For mode HR, the confidence cutoff is set to the value for a 95% yield on the validation set. The respective performances per dataset are reported in Suppl. Table SN.2.7.

| Model | Dataset | Reads classified as noise [%] | mode | Confidence cutoff | Unclassfied reads [%] | Accuracy | Precision | Recall |
|---|---|---|---|---|---|---|---|---|
| WDX-DPC | 1 | 0.0000 | HR | 0.1120 | 1.9000 | 0.9400 | 0.9250 | 0.9450 |
| WDX-DPC | 1 | 0.0000 | HA | 0.2250 | 4.1000 | 0.9460 | 0.9330 | 0.9500 |
| WDX-DPC | 2 | 0.0000 | HR | 0.1120 | 1.8000 | 0.9450 | 0.9380 | 0.9290 |
| WDX-DPC | 2 | 0.0000 | HA | 0.2250 | 3.0000 | 0.9510 | 0.9440 | 0.9370 |
| WDX4 | 3-5 | 1.7500 | HR | 0.7700 | 5.5000 | 0.9924 | 0.9923 | 0.9924 |
| WDX4 | 3-5 | 1.7500 | HA | 0.9400 | 11.0400 | 0.9943 | 0.9941 | 0.9943 |
| WDX4 | 6 | 4.6700 | HR | 0.7700 | 9.1900 | 0.9899 | 0.9900 | 0.9864 |
| WDX4 | 6 | 4.6700 | HA | 0.9400 | 16.9200 | 0.9926 | 0.9925 | 0.9917 |
| WDX6 | 3-5 | 2.1900 | HR | 0.6600 | 4.6900 | 0.9949 | 0.9949 | 0.9949 |
| WDX6 | 3-5 | 2.1900 | HA | 0.8900 | 10.5000 | 0.9968 | 0.9969 | 0.9968 |
| WDX6 | 6 | 5.7300 | HR | 0.6600 | 8.6900 | 0.9921 | 0.9925 | 0.9899 |
| WDX6 | 6 | 5.7300 | HA | 0.8900 | 16.2700 | 0.9952 | 0.9957 | 0.9940 |
| WDX8 | 3-5 | 2.5600 | HR | 0.5600 | 4.7500 | 0.9912 | 0.9914 | 0.9911 |
| WDX8 | 3-5 | 2.5600 | HA | 0.8200 | 9.4200 | 0.9948 | 0.9948 | 0.9947 |
| WDX8 | 6 | 5.6000 | HR | 0.5600 | 7.9800 | 0.9851 | 0.9853 | 0.9834 |
| WDX8 | 6 | 5.6000 | HA | 0.8200 | 14.6800 | 0.9900 | 0.9902 | 0.9897 |
| WDX10 | 3-5 | 2.5200 | HR | 0.4835 | 5.2700 | 0.9870 | 0.9871 | 0.9869 |
| WDX10 | 3-5 | 2.5200 | HA | 0.7370 | 10.0200 | 0.9924 | 0.9925 | 0.9922 |
| WDX10 | 6 | 5.3100 | HR | 0.4835 | 8.5700 | 0.9827 | 0.9818 | 0.9813 |
| WDX10 | 6 | 5.3100 | HA | 0.7370 | 14.6500 | 0.9890 | 0.9886 | 0.9877 |
| WDX12 | 3-5 | 2.1700 | HR | 0.4315 | 5.5100 | 0.9788 | 0.9791 | 0.9785 |
| WDX12 | 3-5 | 2.1700 | HA | 0.7200 | 10.9300 | 0.9853 | 0.9855 | 0.9850 |
| WDX12 | 6 | 4.7500 | HR | 0.4315 | 8.8500 | 0.9717 | 0.9697 | 0.9717 |
| WDX12 | 6 | 4.7500 | HA | 0.7200 | 15.7000 | 0.9811 | 0.9798 | 0.9805 |

**Table SN.2.7:** Performance Comparison Across Models in Different Operation Modes

#### Performance of individual barcodes in different barcode sets

##### WDX4

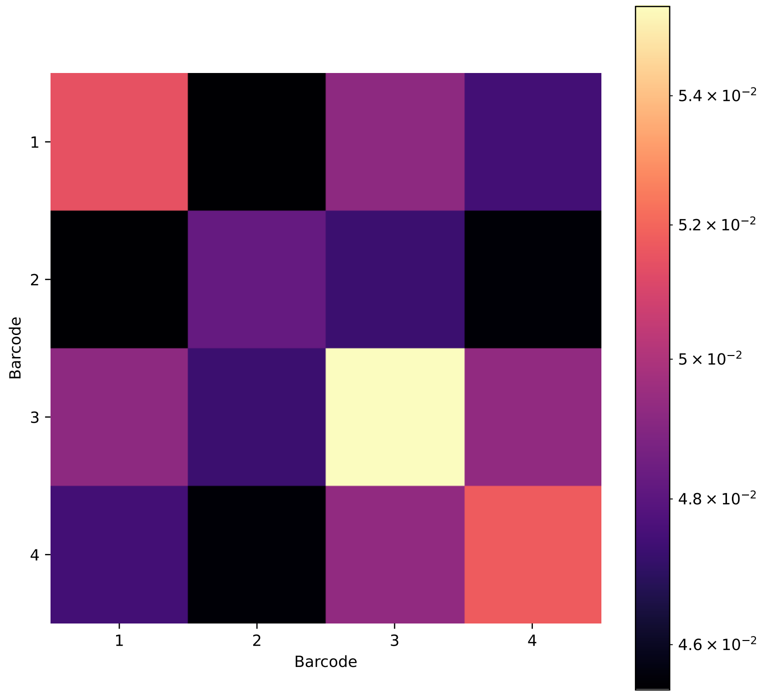


**Figure SN.2.2:** The kernel matrix used to train the WDX4 model, with values averaged per class pair, shows the separation between barcode classes.

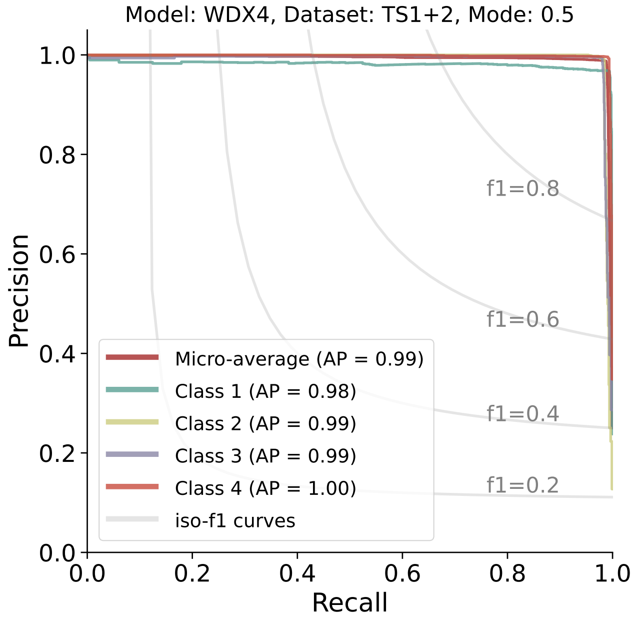


**Figure SN.2.3:** Precision-recall curves of individual WDX-RTA barcodes used in WDX4 model. Evaluated on dataset 3–6 at a confidence cutoff of 0.5.

#### WDX12

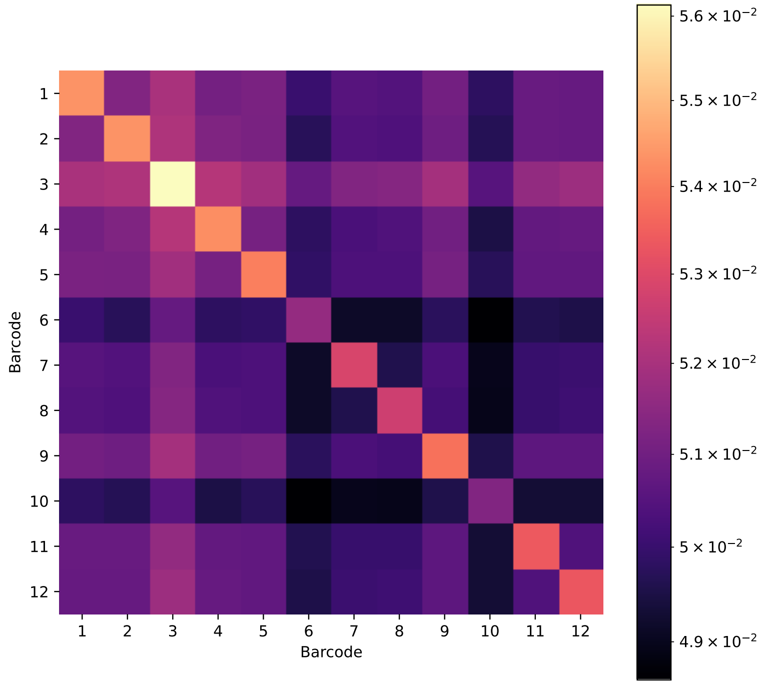


**Figure SN.2.4:** The kernel matrix used to train the WDX12 model, with values averaged per class pair, shows the separation between barcode classes.

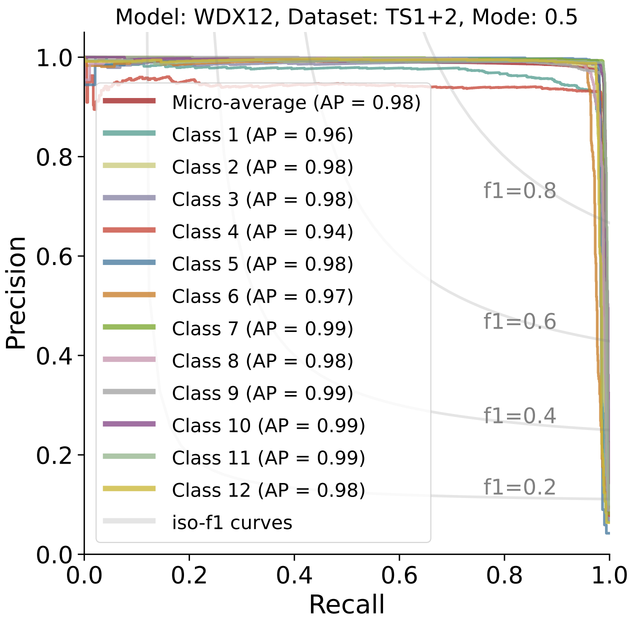


**Figure SN.2.5:** Precision-recall curves of individual WDX-RTA barcodes used in WDX12 model. Evaluated on dataset 3–6 at a confidence cutoff of 0.5.

**DPC-RTA**

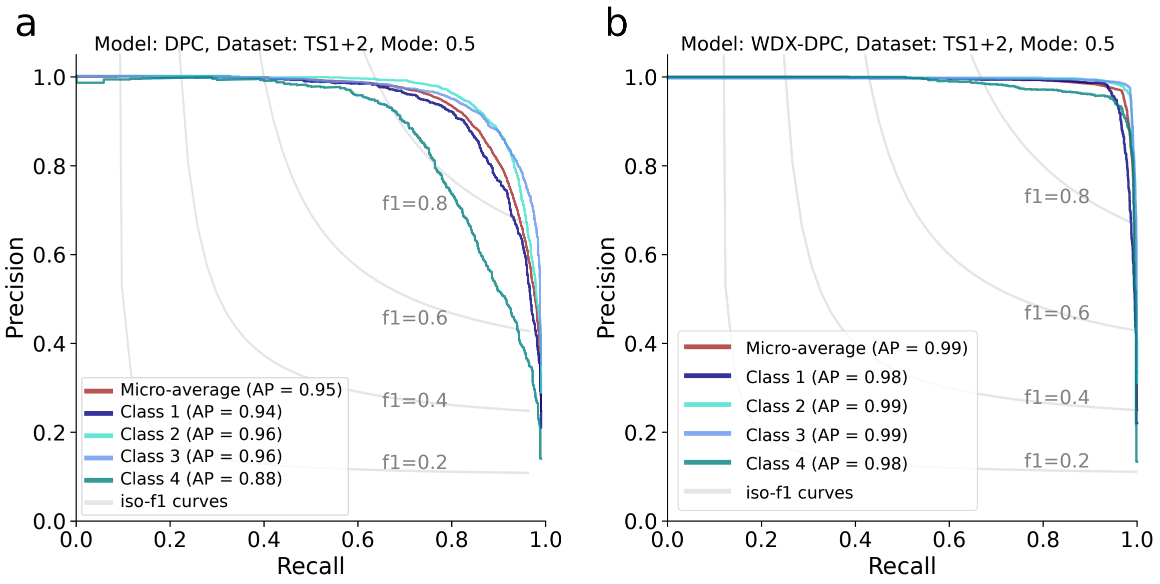


**Figure SN.2.6:** Precision-recall curves for DeePlexiCon (left; (a)) and WDX-DPC (right; (b)) model. Evaluated on dataset 1–2 at a confidence cutoff of 0.5. Both models were trained on the same data (provided by the DeePlexiCon authors), and both evaluated on datasets 1 and 2. Left: Performance of published DeePlexiCon model. Right: Performance of WarpDemuX model.

#### Short poly(A) tail length

Notably, WarpDemuX signal preprocessing hinges on accurately detecting the (start of the) polyA tail in the raw signal, which becomes increasingly challenging as the polyA tail length decreases. Despite this, the tool also achieved high demultiplexing performance on RNA with short polyA tails (as in dataset 6). This indicates that WarpDemuX can also be employed in experimental settings where enzymatic poly-adenylation is not feasible, such as in the studies were the length of the polyA tail is of interest [41, 42, 43, 3].

#### SN.3 Analysis of SARS-CoV-2 sequencing run

Vero cells were infected with recombinant SARS-CoV-2 wildtype or SARS-CoV-2-TurboGFP, replacing ORF8. This results in distinct subgenomic RNAs being expressed by either variant SN.3.1.

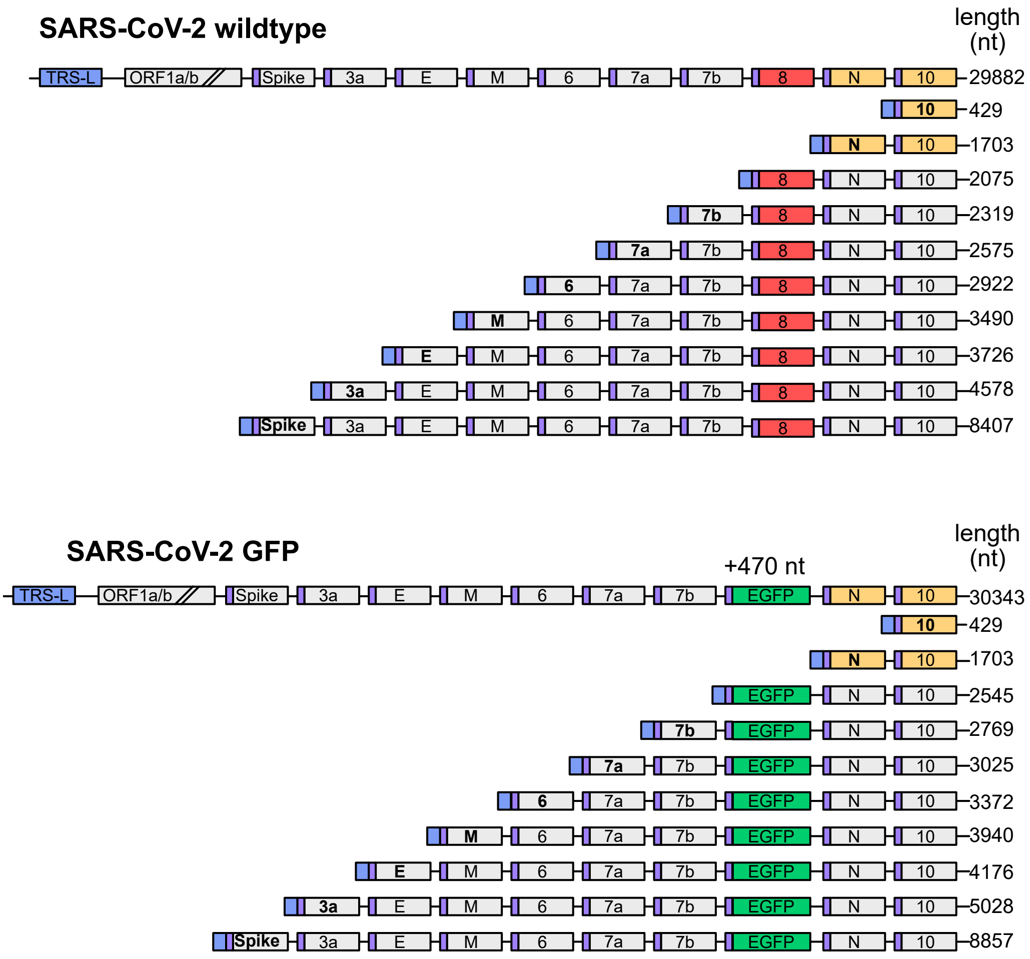


**Figure SN.3.1:** Schematic of viral RNAs generated during infection with the SARS2-CoV-2 wildtype (upper) or GFP-expressing variant (lower). TRS-L sequence is marked in light blue, the expressed isoform for subgenomic RNAs is boldened. The two isoforms that are not distinguishable between the wildtype and GFP variant are colored in yellow.

Cellular RNA and supernatant RNA was extracted using phenol/chlorofom and barcoded with WDX barcodes as described in SN.3.1.

| Sequencing run | WDX barcode | Sample | Location |
|---|---|---|---|
| 5 | WDX_bc01 | Vero-TMPRSS2 mock | cell |
| | WDX_bc02 | Vero-TMPRSS2 inf. with SARS-CoV-2 8 h p.i. | cell |
| | WDX_bc03 | Vero-TMPRSS2 inf. with SARS-CoV-2 24 h p.i. | cell |
| | WDX_bc04 | Vero-TMPRSS2 inf. with SARS-CoV-2 48 h p.i.m | cell |
| | WDX_bc05 | Vero-TMPRSS2 inf. with SARS-CoV-2 72 h p.i. | cell |
| | WDX_bc06 | Vero-TMPRSS2 inf. with SARS-CoV-2 72 h p.i. | supernatant |
| | WDX_bc07 | Vero-TMPRSS2 mock | cell |
| | WDX_bc08 | Vero-TMPRSS2 inf. with SARS-CoV-2-GFP 8 h p.i. | cell |
| | WDX_bc09 | Vero-TMPRSS2 inf. with SARS-CoV-2-GFP 24 h p.i. | cell |
| | WDX_bc10 | Vero-TMPRSS2 inf. with SARS-CoV-2-GFP 48 h p.i. | cell |
| | WDX_bc11 | Vero-TMPRSS2 inf. with SARS-CoV-2-GFP 72 h p.i. | cell |
| | WDX_bc12 | Vero-TMPRSS2 inf. with SARS-CoV-2-GFP 72 h p.i. | supernatant |

**Table SN.3.1:** WarpDemuX RTA adapter to sample assignment of SARS-CoV-2 sequencing run

#### Run statistics

After sequencing, reads were filtered for a mean Phred quality score of 10 (Sup. Fig. SN.3.2a). All reads were then passed through the WarpDemuX pipeline. Notably, we observed a relation between the basecall-based read Phred qscore and the WDX confidence score distribution (Sup. Fig. SN.3.2b-e), indicating that there is a common underlying source of noise that can affect both basecalling and WDX demultiplexing confidence [44].

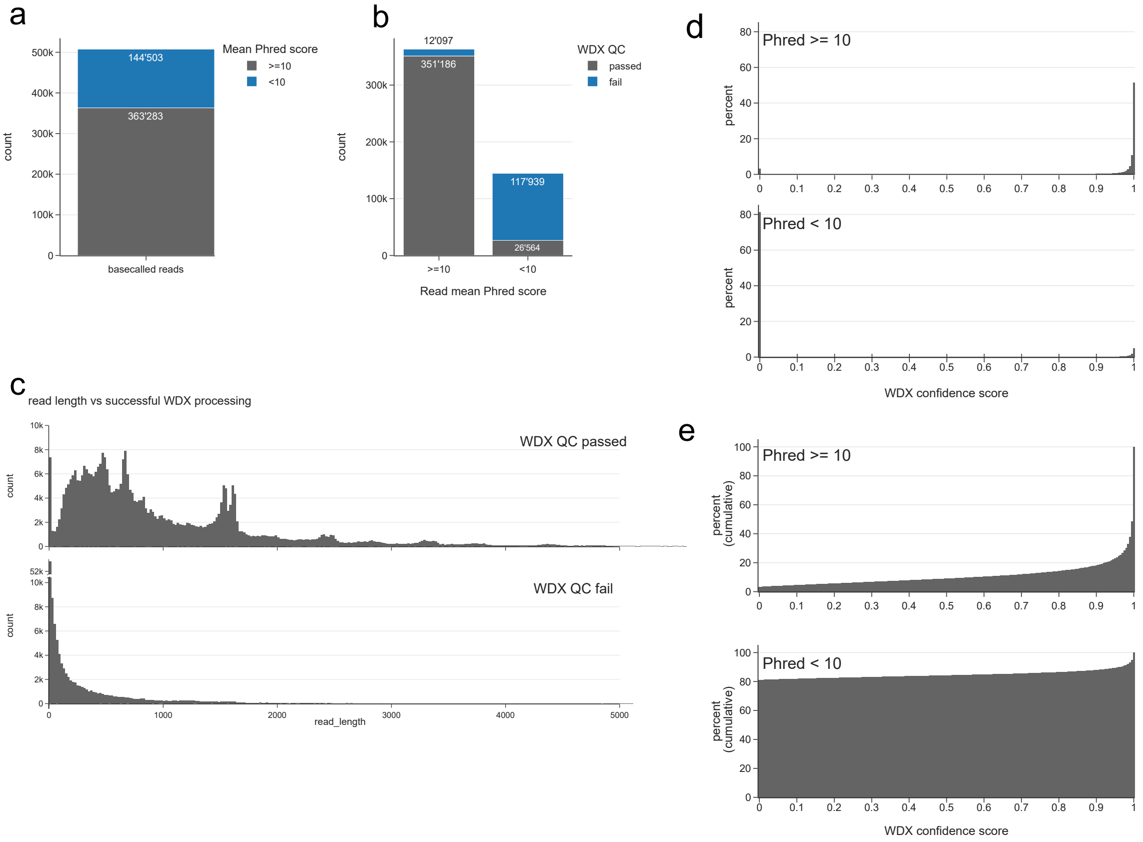


**Figure SN.3.2:** Run statistics for the multiplexed SARS-CoV-2 sequencing run. (a) Total number of basecalled reads, split by mean read Phred score 10. (b) Number of reads passing WarpDemuX quality checks (poly(A) detection, signal quality checks) separated by mean read Phred qscore of 10. (c) Read length distribution of reads passing or failing WarpDemuX QC. (d) WarpDemux confidence score distribution for reads passing WDX QC separated by read mean qscore of 10. (e) Cumulative WarpDemuX confidence score distribution for reads passing WDX QC separated by read mean qscore of 10.

#### Read to species assignments

Next, reads were aligned with minimap2 to the Vero reference genome (Genbank: GCA_000409795.2), SARS-CoV-2 genome (NCBI RefSeq: LC528233) or TurboGFP sequence (NCBI RefSeq: GU452685.1 306 to 999):

```
minimap2 -ax splice -u f -k14 {reference_index} {fastq_file} > {bamfile}
```

Uniquely mapped reads to each reference were extracted with samtools:

```
samtools view -F 0x100 {bamfile} | awk '{{print $1, $3}}'  > {outfile}
```

When a multi-mapping read was detected it was assigned to $ref_A$ - $ref_B$ (for example reads that aligned to both SARS-CoV-2 and TurboGFP were assigned to SARS2-TurboGFP). Those aligning to multiple Vero reference contigs were assigned "Vero". Already at a WarpDemuX confidence threshold of 0 the vast majority of reads classified as a given barcode were of the expected species origin. Only for the two highly underrepresented barcodes 6 and 12 (supernatant samples) some SARS2-GFP reads were detected, which are likely from a low level of spillover from other samples that becomes substantial with low number of true reads.

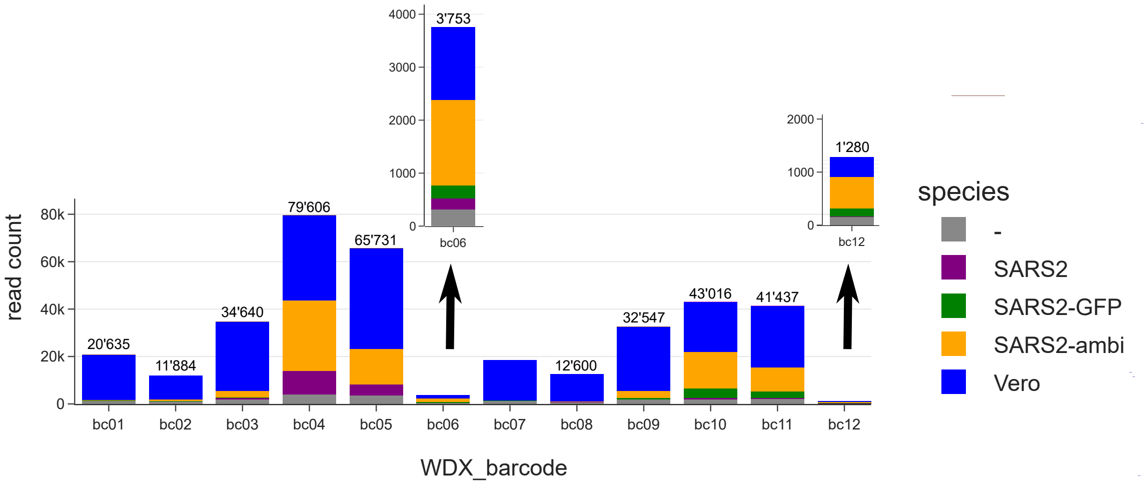


**Figure SN.3.3:** Run statistics after WDX demultiplexing (without WDX confidence threshold). Number of reads classified per barcode at a WDX confidence score cutoff of 0, colored by species. The numbers describe the total read count per barcode.

#### WarpDemuX confidence threshold choice

The WarpDemuX confidence score represents the confidence of the classifier to correctly classify a given adapter signal. The confidence threshold below which reads are assigned as unclassified can be chosen depending on the experimental setup and requirements and results in a tradeoff between yield and accuracy (SN.2.4). In this case, for heavily unbalanced input containing two rare (supernatant) samples we determined that a highly stringent cutoff must be chosen to ensure high specificity even for these rare samples. Upon evaluating the confidence score distributions per sample (Sup. Fig. SN.3.4) a confidence threshold of .99 was chosen to optimally balance demultiplexing accuracy with yield. While this resulted in approximately 32% unclassified reads across all samples (Sup. Fig. SN.3.5a), the fraction of reads below a WarpDemuX confidence score of 0.99 was substantially higher for the two rare samples bc6 and 12 (80% and 64% respectively) (Sup. Fig. SN.3.5b), consistent with higher relative levels of spillover expected. Indeed, with this stringent threshold almost no Vero reads were now detected in the supernatant samples, as well as no GFP reads remaining in the sample with bc06 (Sup. Fig. SN.3.5c).

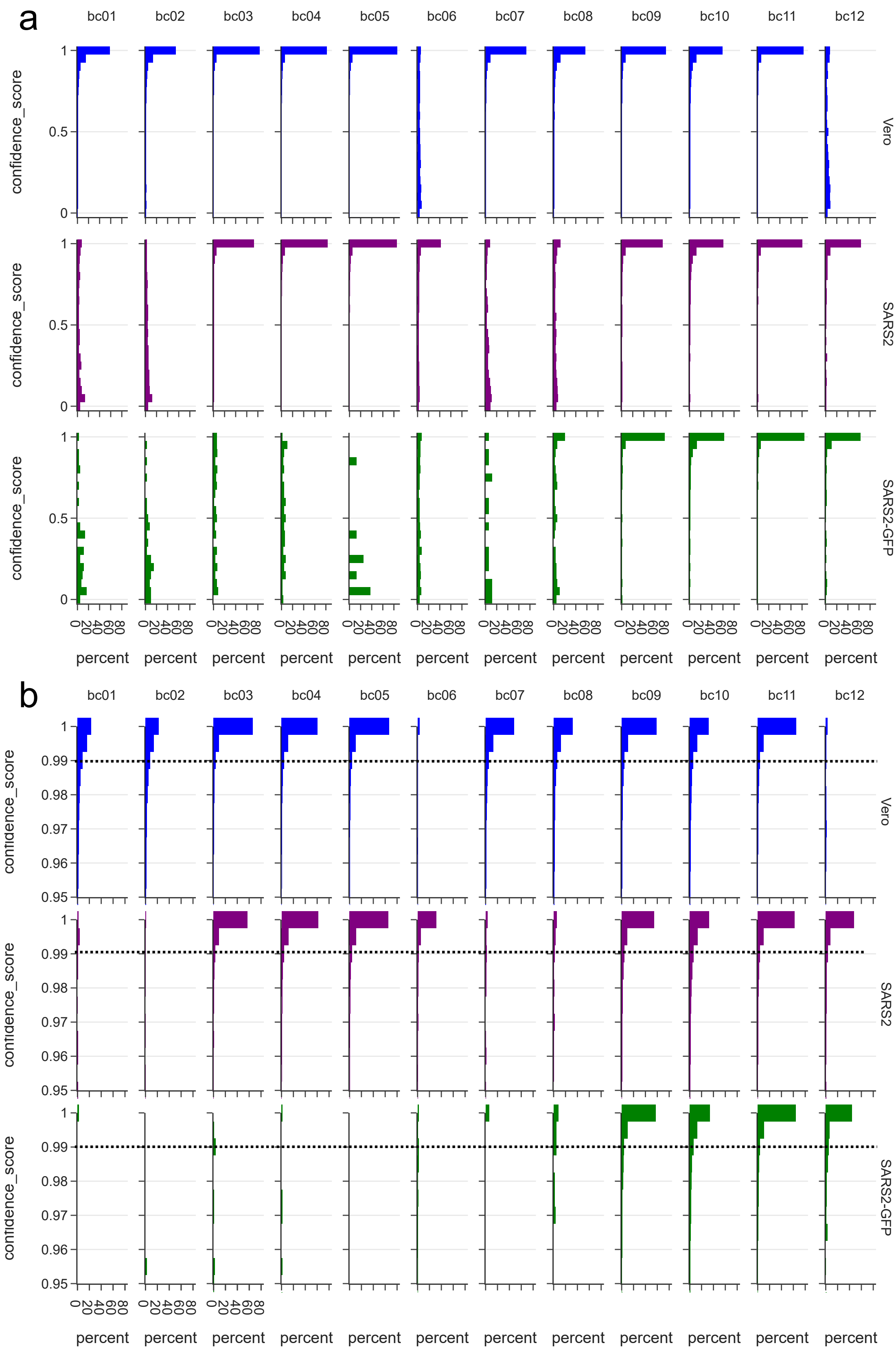


**Figure SN.3.4:** WarpDemuX confidence threshold distribution per barcode and species. (a) WarpDemuX confidence score distribution (normalized as percent) per barcode (column) and species (row). (b) Zoom of (a) with higher resolution for confidence scores between 0.95 and 1.

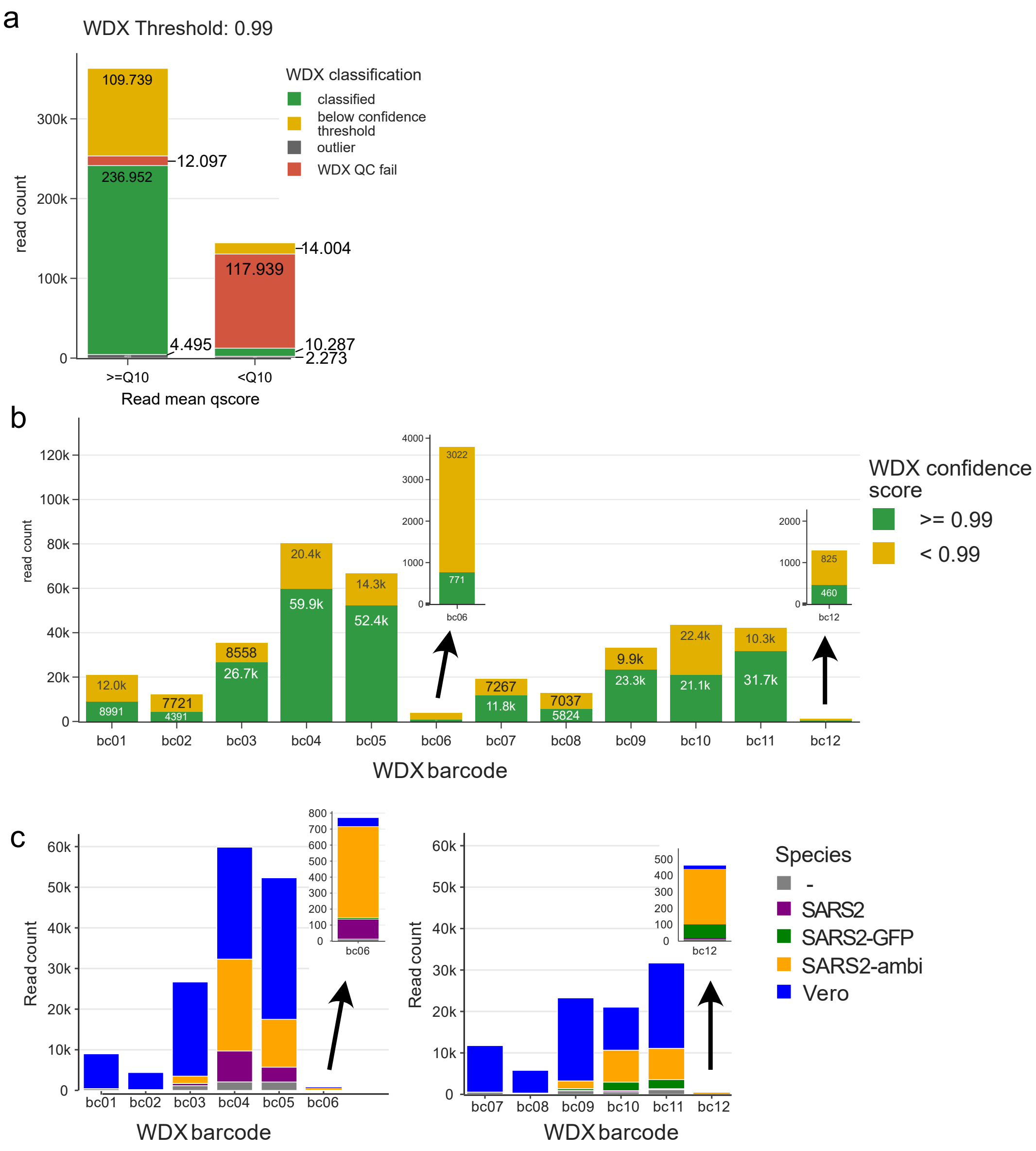


**Figure SN.3.5:** WarpDemuX confidence threshold of 0.99. (a) Number of reads passing a WarpDemuX confidence threshold of 0.99 by read mean Phred score. (b) Number of reads per barcode passing the 0.99 confidence threshold. (c) Total number of reads aligning uniquely to the genome of a species per barcode at a WDX confidence threshold of 0.99.

When evaluating the read lengths after WarpDemuX demultiplexing through a virtual gel, bands presumably corresponding to SARS-CoV-2 isoforms became immediately apparent (Sup. Fig. SN.3.1). For samples with the GFP variant, the bands above 1.5 kb were shifted approximately 500 bp upwards, consistent with the expected increased length of sgRNAs due to the GFP insert (Sup. Fig. SN.3.1).

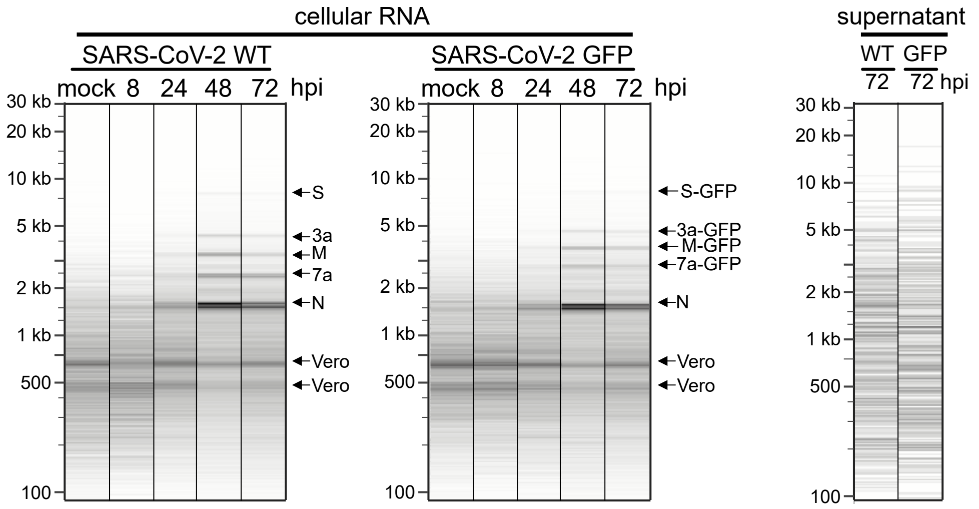


**Figure SN.3.6:** Virtual gel of demultiplexed samples. Linear length scale, normalized per sample. The annotations highlight bands corresponding to SARS-CoV-2 isoforms, with the GFP variant offset by around 500 nt above the WT variant.

#### SARS-CoV-2 read transcript assignments

To confirm that these reads with distinct lengths were full length SARS-CoV-2 subgenomic RNAs, their TRS-L sequence was searched for using cutadapt v4.8 was used with the following settings:

```
SARS2_5UTR="ATTAAAGGTTTATACCTTCCCAGGTAACAAACCAACCAACTTTCGATCTCTTGTAGATCTGTTCTCTA"

cutadapt -g $SARS2_5UTR -e 0.25 --overlap 10 --action=none --untrimmed-output
```

Reads where the TRS-L sequence was found were marked as full-length, while those without TRS-L sequence were labeled as incomplete. When evaluating read lengths of the so classified reads, indeed reads where the TRS-L sequence was present showed distinct lengths SN.3.7, which was subsequently used for isoform assignment by length (Sup. Fig. SN.3.1). If the reads were too short to be distinguished between the GFP and non-GFP variant of the virus, they were assigned as ambiguous.

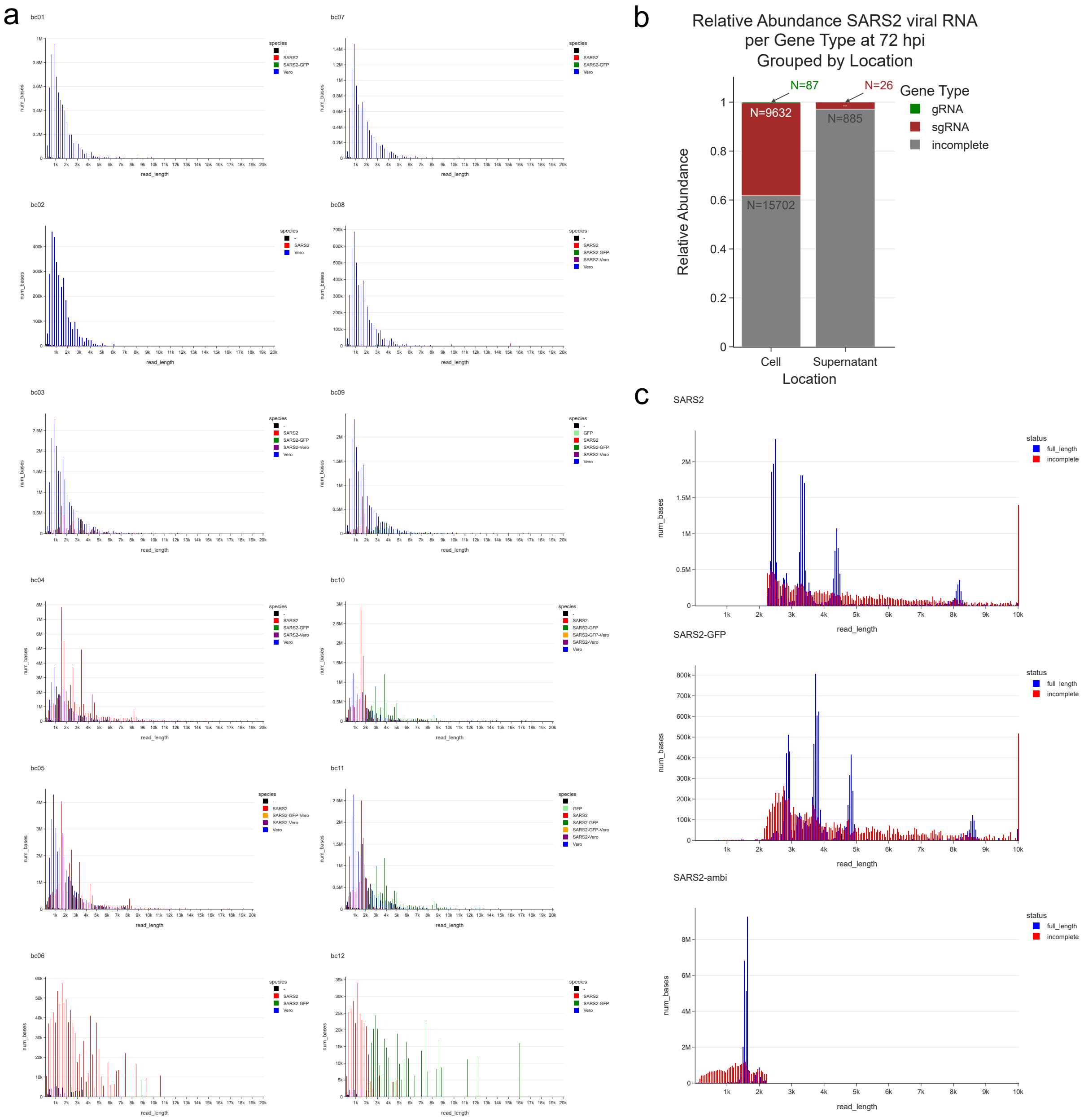


**Figure SN.3.7:** Read length distribution per barcode and species. Classification of SARS2 reads according to presence of TRS-L sequence. (a) Histogram of number of bases by read length for reads mapping to the different species (not split by full length). (b) Relative abundance of SARS-CoV-2 reads for which a TRS-L sequence was detected separated by location of RNA. (c) Read length distributions for full length and incomplete reads aligning to either SARS2-WT (top), SARS2-GFP (middle) or ambiguously to SARS2 (bottom).

After assignment of SARS-CoV-2 isoforms, the relative abundance of full length and incomplete SARS-CoV-2 reads per sample could be estimated, revealing a constant ratio of full length to incomplete reads for the cellular samples throughout infection, while the two supernatant samples showed distinct reduction of full length reads (Sup. Fig. SN.3.8), consistent with the assumption that subgenomic RNA is selected against during packaging.

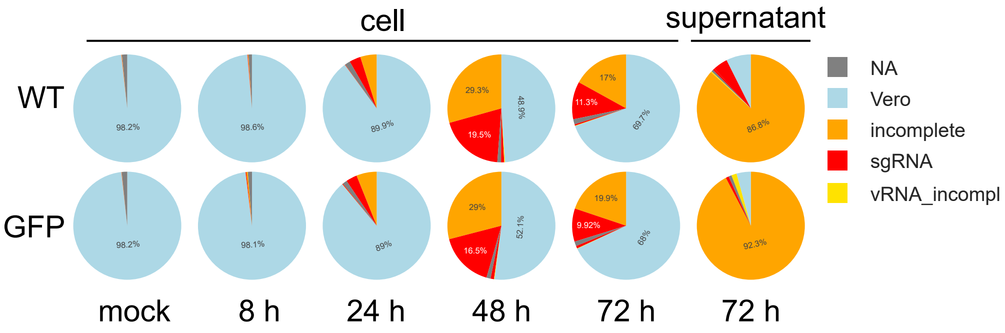


**Figure SN.3.8:** Pie charts of relative abundance of reads aligning to Vero and SARS2, split by incomplete and full length transcripts for SARS2.

#### Vero read transcript assignments

To differentiate RNA types within Vero-aligned reads, we performed read classification with Isoquant v3.3.1. We used transcript annotations contained in the Genbank entryy GCA_000409795.2, as these resulted in higher number of classified reads in comparison the more recent reference GCA_015252025.1. The specific settings used were:

```
isoquant.py --data_type nanopore --stranded forward --complete_genedb --reference {fasta}
        --genedb {gtf} --bam {' '.join(bam_files)} --labels {' '.join(samples)} -o {outdir}
```

Finally, descriptive gene and transcript type annotations were acquired through Ensembl BioMart and merged into the isoquant read classification. This revealed that the vast majority of Vero-aligning reads were from protein-coding genes, as expected from sequencing poly-adenylated RNA. A minor fraction of reads were assigned to mitochondrial rRNA however, possibly due to mitochondrial poly-adenylation of non-coding RNA (see Sup. Fig. SN.3.9a).

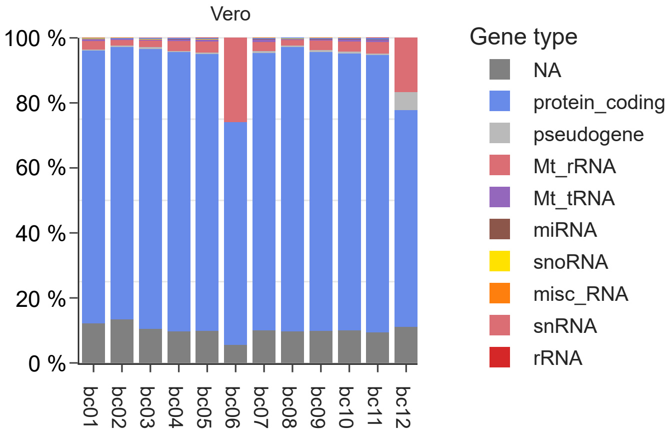


**Figure SN.3.9:** Relative bar charts per sample of gene assignments (generated by isoquant) for reads aligning to the Vero genome.

#### Poly(A) tail length estimation

To estimate the poly(A) tail lengths of SARS-CoV-2 RNA in cells and virion as well as the effect of SARS-CoV-2 infection on cellular mRNA poly(A) tail length we performed tail length estimation with tailfindr [43]. Specifically, we converted the raw pod5 files into fast5 files before rebasecalling with guppy 6.3.4 with ‘–fast5-out‘ option. We then ran tailfindr version 1.4 on the generated fast5 files and incorporated the per read poly(A) tail estimations as provided in the csv file into our analysis.

By leveraging annotation information of Isoquant-assigned transcripts we were also able to study the poly-adenylation landscape of Vero-derived transcripts. We detect distinct poly(A) tail length distributions between nuclear (50-150 nt) and mitochondrial (10-50 nt) transcripts (Sup. Fig. SN.3.10a), as well as distinct mean length differences between SARS2- and Vero-derived transcripts (Sup. Fig. SN.3.10b). Intriguingly, as highlighted in the results, a distinct bimodal distribution of poly(A) tail lengths was apparent for SARS2 transcripts at 72 hours post infection (hpi), with the smaller tail length population being less abundant at earlier time points. A bimodal Gaussian Fit revealed the modes of these two distributions to be 52 nt and 29 nt respectively (Sup. Fig. SN.3.10c), with components $\mathcal{N}_1(29.44, 5.89)$ and $\mathcal{N}_2(51.80, 20.17)$ and mixing coefficients 0.148 and 0.842, respectively. Leveraging our longitudinal samples, we examined the dynamics of the poly(A) tail length in SARS2 cellular transcripts. We then applied the expectation-maximization (EM) algorithm to estimate the mixing coefficients for the 24 and 48 hpi data, while keeping the distribution components fixed. This yielded coefficients of 0.001 and 0.999 for 24 hpi, and 0.272 and 0.973 for 48 hpi.

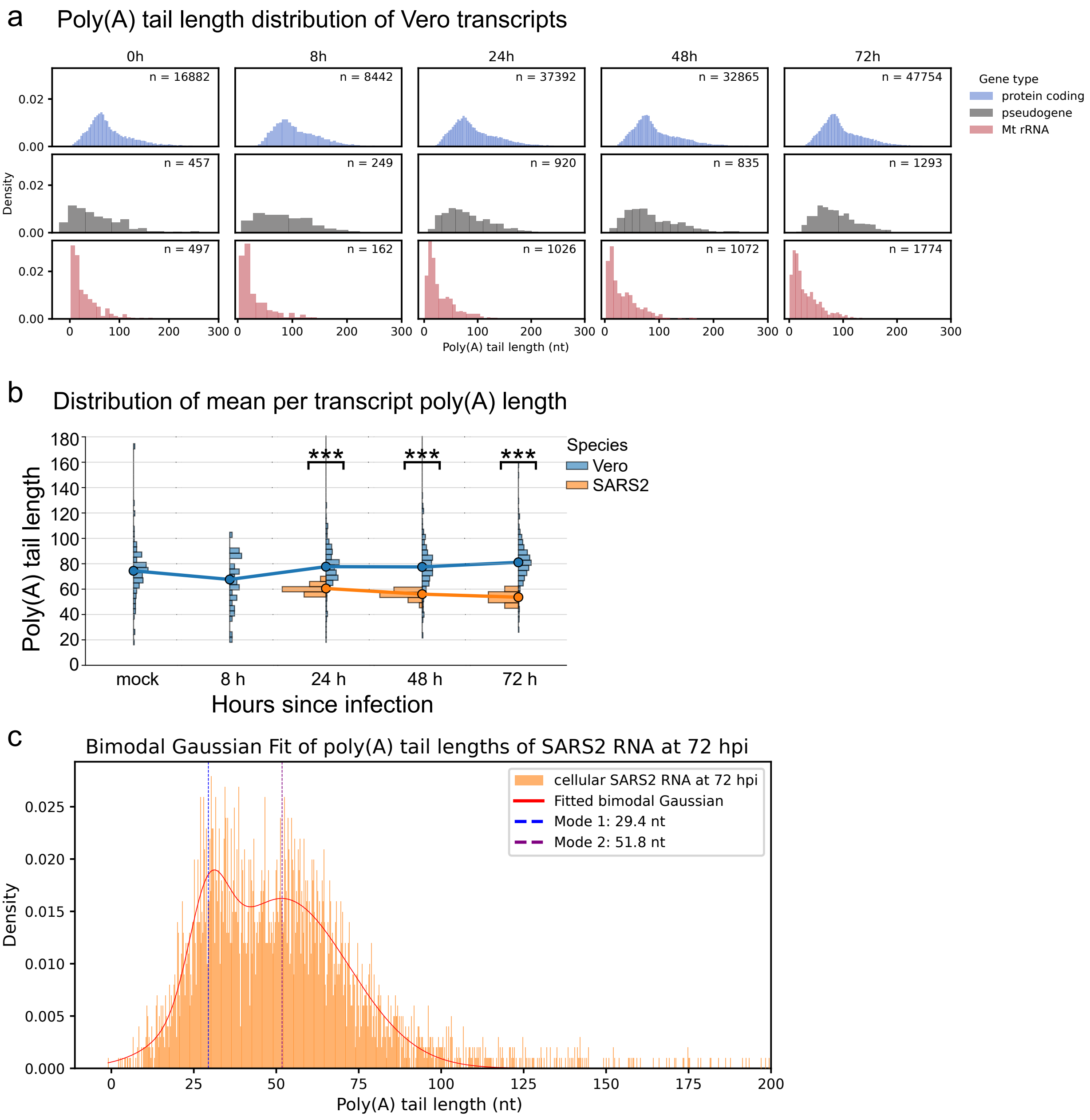


**Figure SN.3.10:** Poly(A) tail length distribution statistics. (a) Poly(A) tail length distributions of Vero transcripts by gene type (biomart annotation). (b) Poly(A) tail length distribution per transcript. For each transcript with coverage of at least 20 the mean poly(A) tail length was calculated. (c) Bimodal Gaussian fit of the poly(A) tail length distribution of SARS2 reads at 72 hpi to determine the modal poly(A) tail length for the two distributions.

### SN.4 Live barcode balancing

#### Latency of WarpDemuX adaptive-sampling

WarpDemuX's classification speed allowed the implementation of barcode-specific adaptive sampling for dRNA-seq. Our goal was to make rejection decisions while the poly(A) tail of RNA molecules is still within the pore such that i) rejected reads do not end up in the sequencer output folder (`fast5_pass`/`pod5_pass` and ii) the risk of pore blockage upon rejection is reduced by preempting the formation of RNA structures on the trans side of the pore that may hinder ejection of the molecule back through the pore.

Indeed, when evaluating the rejected reads after completion of the adaptive-sampling run, most rejected reads were too short to be written out to `pod5` files (**Fig. SN.4.1a**), with rejection request successfully executed within 400 ms after detection of the polyA tail for nearly all reads (**Fig. SN.4.1b**). Reads that were rejected but still written out to **pod5** files exhibited shorter read lengths compared to retained reads (**Fig. SN.4.1c**).

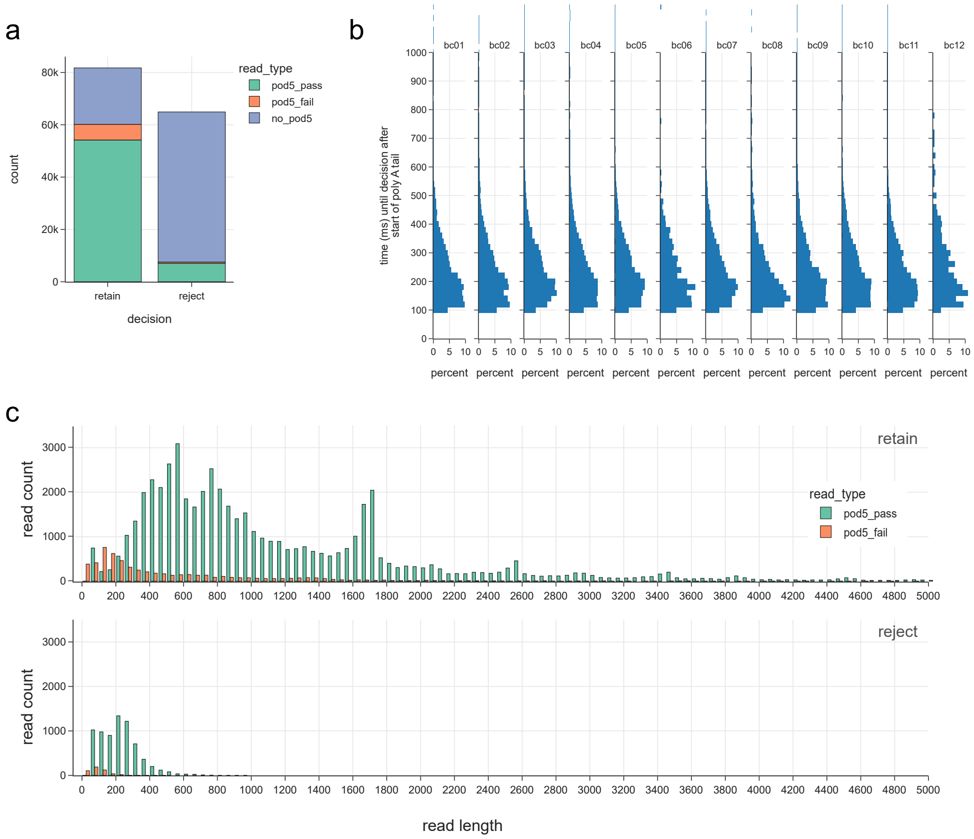


**Figure SN.4.1:** Latency of live WarpDemuX balancing. (a) Reads with reject decisions are most of the time too short to be written into pod5 files. (b) The duration between the start of the poly(A) sequence and the reject decision is constant between barcodes. (c) Read length distribution (after basecalling) for reads that are written into pod5 files, separated by decision and qscore filter.

#### Balancing strategies

Relative barcode abundances can be balanced based on different statistics. We developed multiple balancing strategies, such as barcode balancing based on adapter counts, read counts, or number of bases (**Fig. SN.6**). We also implemented barcode blacklisting, where certain barcodes are automatically rejected, independent of the barcode balance.

To evaluate the performance of the different methods, we applied each to 40 channels each on a MinION R9.4.1 flow cell, leaving 200 channels to sequence without adaptive-sampling for comparison. Balancing strategies resulted in a more even read count over barcodes, and the blacklist logic depleted targeted barcodes only, as expected (**Fig. SN.4.2**).

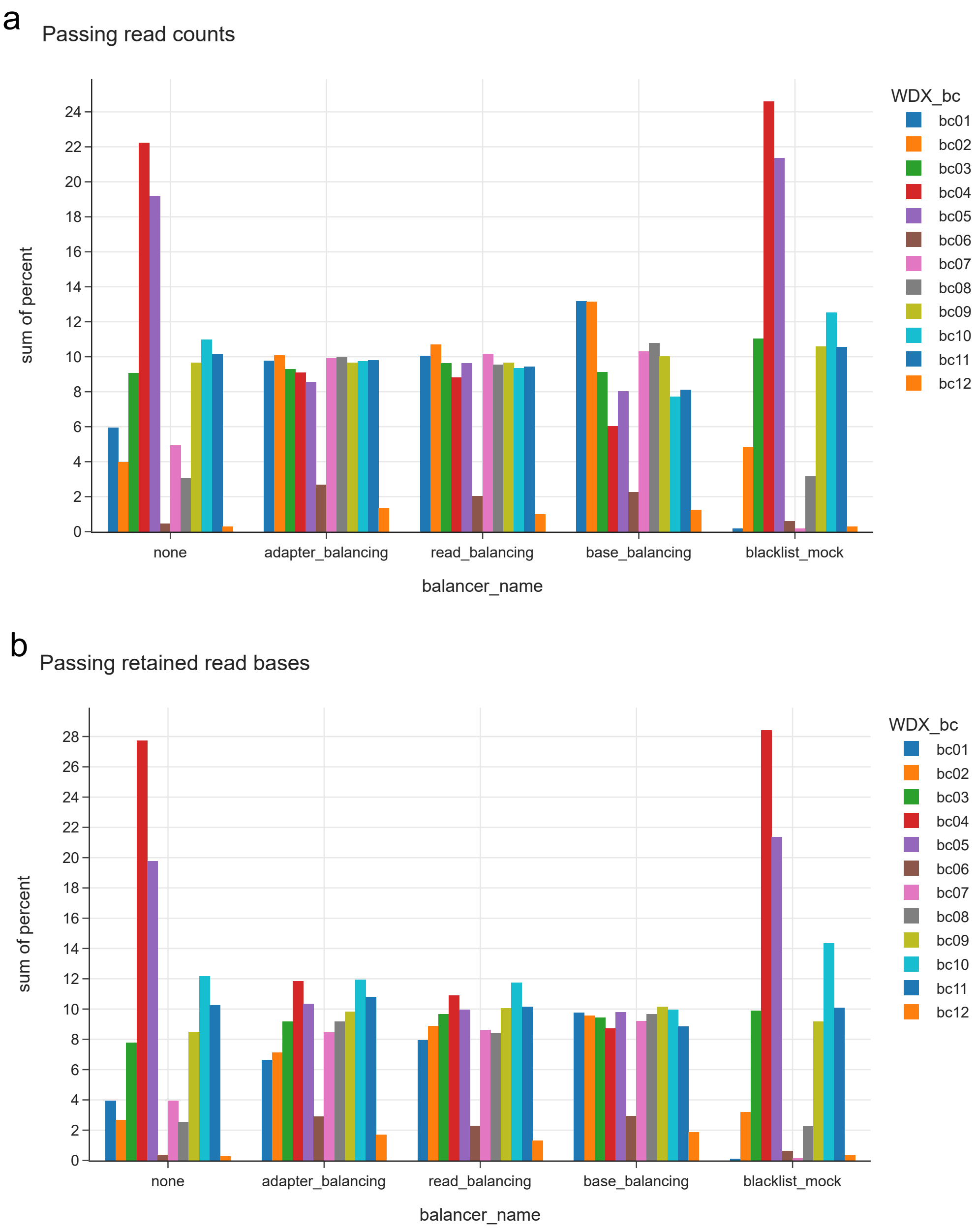


**Figure SN.4.2:** Balancer Strategies. Balancer name 'none' indicates no adaptive-sampling. In 'blacklist_mock', barcode 1 and 7 were blacklisted. (a) Relative abundance of reads for each barcode when different balancer strategies are applied. (b) Relative abundance of bases for each barcode when different balancer strategies are applied.

#### Pore lifetime

Ultra-fast barcode classification with adaptive-sampling can lead to rejection frequencies above 50% (4e), depending on the abundances of the multiplexed samples and balancing strategy. An unsuccessful rejection may result in pore blockage, reducing the lifetime of the pore and sequencing yield of the run. We aimed to minimize the number of pore blockages upon unsuccessful rejection by rejecting reads while they are still within the poly(A) tail. We analyzed the effect of different balancing strategies on pore lifetime, and found no substantial differences between pores with and without adaptive sampling, regardless of the strategy used (**Fig. SN.4.3**).

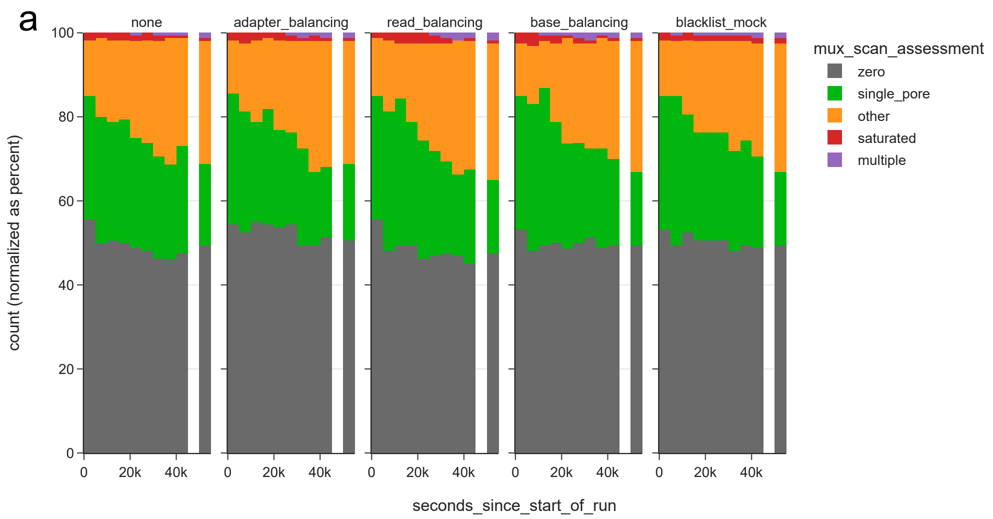


**Figure SN.4.3:** Pore lifetime with different balancer strategies. Visualization of pore scan results during the live adaptive sampling sequencing run.

#### SN.5 Temporal Perturbation Analysis between DTWD and GASF

While the DTWD is resilient to temporal perturbations, the GASF feature transformation exhibits vulnerability to such perturbations (Fig. SN.5.1). Consequently, stochastic fluctuations in dwell time introduce significant noise when employing a GASF-based approach. Although DeePlexiCon shows that a CNN can learn to discern between genuine correlations and noise in the signal-transformed image, the inherent complications of the GASF transformation intensify the classification task. This is manifested in the augmented training data requisite for the CNN and its diminished performance in comparison with WarpDemuX.

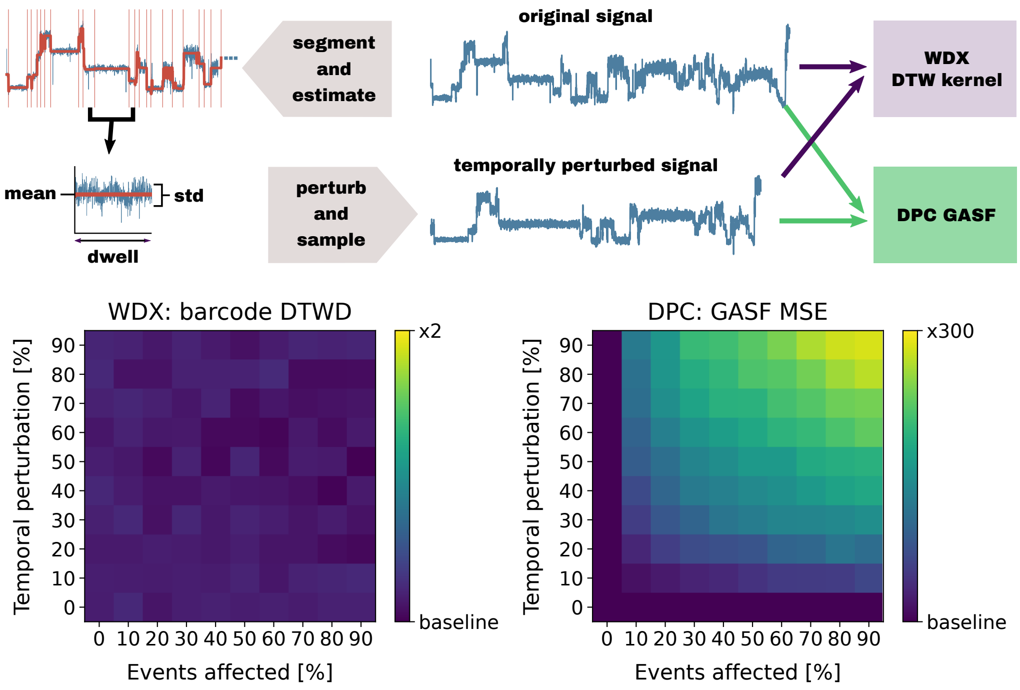


**Figure SN.5.1:** Temporal Perturbation Analysis between DTWD and GASF. A comparative study illustrating the susceptibility of the GASF feature transformation to temporal perturbations, as opposed to the robustness exhibited by DTWD.

### SN.6 Appendix: Software Adaptations

Live barcode balancing in direct RNA sequencing has the prospect to substantially increase yield for multiplexed sequencing runs, as well as to increase pore half-life when some, but not all samples that are sequenced have a high likelihood of blocking pores. It therefore can reduce labour and resource costs, making dRNA-seq a more viable solution to prevalent questions related to epitranscriptomics.

Of note is that adaptive barcode balancing will not alter the relative read abundance of barcodes, but it will result in higher turnover of reads, thus leading to a decreased of sequenced bases for rejected and concurrent increase of bases for accepted barcodes.

To facilitate live barcode balancing, multiple steps need to be performed with high accuracy, yield and speed: 1.) streaming data quickly from the sequencing device, 2.) identifying and extracting adapter signals from the start of a read until the transition to polyA tail/RNA, 3.) rapid processing and classification of the adapter signal, and lastly, 4.) decision on rejection and if so, sending the command to the sequencer.

For dRNA adaptive barcode sampling to result in substantial enrichment when compared to a standard sequencing run, the time from the read starting to enter the pore to the decision should be as short as possible. To estimate theoretical limits of enrichment, we can Additionally, rejection of RNA reads should not lead to a substantial increase in pore loss.

One possible source of pore loss can be the RNA refolding at the trans side of the pore. To reduce the likelihood of this, we aim to reject reads while the poly(A) tail is still in the pore. To estimate the required turn-around time for this goal for RNA002 we calculate the following: DNA adapter translocation takes approximately 2 seconds, followed by translocation of RNA nucleotides at 70 bases per second. At a poly(A) tail length of 30 nucleotides, this means optimally we want to achieve a time of up to 430 ms from poly(A) start to decision.

The adaptive sampling process is accessible through the read_until_api, which acts as an interface between the gRPC interface in MinKNOW and custom python code. Matt Loose's team leveraged this API to develop the first iteration of DNA adaptive sampling [16], adding a critical component, an accumulating cache, to the read_until_api. This cache is critical for us because we require the near whole adapter signal at once (up until the poly(A) signal starts) to be able to segment and classify properly, and this signal is highly likely to stretch multiple individual chunks.

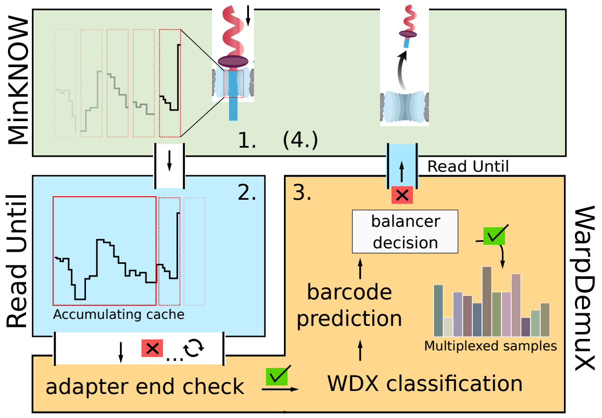


**Figure SN.6.1:** Overview software design WarpDemuX adaptive sampling. 1.) The sequencer (MinKNOW) streams data in chunks of a fixed size. 2.) ReadUntil communicates with the sequencer API and accumulates the read data in an 'Accumulating cache'. 3.) WarpDemuX watches the accumulating cache for the start of a poly(A) ('adapter end check'). When found, the adapter signal is send off to the WarpDemuX classification pipeline. Based on the predicted barcode label and the current sequencing distribution over multiplexed samples, a balancing decision is made (retain/reject). To reject a read, WarpDemuX sends an 'unblock' request to MinKNONW, via ReadUntil.

#### Reducing the Chunk Size

Chunks are blocks of signal that are received through the `read_until_api`. The size of chunks is determined in the sequencing protocol toml file, specifically by the `break_reads_second` parameter. By default chunks consist of 1 second of signal. However, as we attempt to achieve low latency, reducing the chunk size was a critical step. Indeed, reducing the `break_read_seconds` parameter to 0.1 second resulted in receiving chunks of size 300 samples, enabling rejection during or shortly after the poly(A) tail has passed the pore.

However, this change also resulted in nearly no reads being generated by MinKNOW. Investigating this issue further, we identified the `read_classification` parameters as well as the `channel_states` as the two key steps that required tuning. The reason is as follows: first, new reads are detected by an open pore event (which is not bound by the chunk size). Then, each chunk gets classified according to the boundaries specified in the `read_classification` parameters (duration means duration since the read started). The channel state is then determined by regex matching of patterns of chunk classifications, those are defined in the `channel_states.toml` file.

As we decrease the chunk size, statistics such as median and range (90%-10% percentiles) become less consistent, and we identified these as the parameters that required the most tuning. Critically, the range was previously used to detect stalled reads, as it was unlikely (apart from the poly(A) tail) that the signal stayed within a small range for 1 second. In contrast, it may happen relatively frequently that the signal may stay constant in a 0.1 second window–thus we decided to decrease the minimum range required. Upon replaying the simulation, these changes not only recovered the output of the initial sequencing run, but increased it by approximately 25% (data not shown). However, this change in isolation may lead to stalls not being detected, thus severely affecting overall flow cell yield by not recovering blocked pores. To address this problem we added a `read_classification` state "stalled" that chunks where the range of the signal is near 0 should be classified in. Since brief stalls are expected to occur within a read, we tolerate up to 3 'stalled' chunks in between two 'strand' chunks before the `channel_state` is changed from strand to stalled. Only when at least 10 chunks are consecutively classified as 'stalled' do we change the `channel_state` to 'blocked', which will then initiate the standard channel unblocking procedure.

#### Read Until API changes

Having achieved a 100 ms chunk size, we next focused on the read until API. As previously mentioned, the accumulating cache that appends chunk raw signal (instead of overwriting it) is critical for live barcode classification, especially with the decreased the chunk sizes. However, we noticed that for some reads, the start of the signal did not match the start of the new read, and this may result in incorrect poly(A) detection, segmentation or classification. We identified the source of this to stem from a filter on incoming chunks according to their classification implemented in the accumulating cache, and only accepting adapter- or strand-classified chunks. While this is reasonable for basecaller-based adaptive sampling, it can lead to the mentioned data loss at the start of reads due to the possibility of other types of chunk classifications. As such, we have removed this filter during chunk accumulation, but instead added a filter in the `get_read_chunks` function that in our case will only return accumulated chunks that contain at least one adapter classification (see our implementation at `github.com/KleistLab/WarpDemuX`). In addition, we added a max length to the accumulated raw data to ensure that raw signal of long reads or inactive channels does not grow infinitely.

#### Decision Strategies

The last step after classifying raw signal of adapters is to decide on the fate of the read. The simplest approach would be rejection based on relative adapter occurrence. However, this has drawbacks with regards to unligated adapter sequences, which when evaluating raw data may be present in 50% of adapter events, and potentially at varying abundance between different samples.

Instead, we propose two alternative strategies, that both employ additional information present in pod5 files that are generated during the sequencing run. In both strategies we first classify adapters and store their read ids. We then search in the pod5 files as they are generated during the run to identify matching read ids, counting all matches for each barcode. As adapter-only reads are not written out to pod5 files this ensures that we only normalize on true RNA-containing reads. Intriguingly, the output of this analysis can subsequently be used to estimate the fraction of adapter-only reads in the library, helping to troubleshoot library preparation protocols.

In the second strategy we perform the same lookup in pod5 files, but integrate the number of detected events by MinKNOW. These events correspond closely to the number of bases in the read (with approximately 100 additional events introduced by the adapter), so incorporating this information allows us to balance different samples not only by read count, but by base count.

Similarly, if samples are suspected to result in high levels of blockage, we can also identify barcodes that resulted in higher than expected rates of pore blockage by monitoring the end reason of reads in pod5 files, and flag those samples for rejection in order to extend pore lifetime.
